# Cyclic AMP compartmentalization drives signal specificity to control vector colonization and mammalian host infection by American trypanosomes

**DOI:** 10.64898/2025.12.04.692279

**Authors:** Milad Ahmed, Joshua Carlson, Asima Das, Syeda Farjana Hoque, Joshua B. Benoit, Miguel A. Chiurillo, Noelia Lander

## Abstract

Cyclic AMP (cAMP) signaling is crucial for environmental sensing and response to stress conditions in trypanosomatids. However, the mechanisms driving the specificity of cAMP signals remain poorly understood in these protozoan parasites. We recently identified two putative cAMP microdomains in *Trypanosoma cruzi*, the causative agent of Chagas disease. Here, considering the localization of three phosphodiesterases, PDEC at the contractile vacuole complex (CVC), and PDEB1 and PDEB2 along the flagellum, we modulated their expression to functionally characterize the flagellar tip (FT) and the CVC as individual cAMP microdomains, named FT-cAMP and CVC-cAMP, respectively. We generated PDE knockout and overexpression cell lines to selectively alter cAMP signals generated in each compartment. Our results indicate that FT-cAMP mediates cell adhesion, metacyclogenesis, host cell invasion, and intracellular replication, while CVC-cAMP is required for osmoregulation and epimastigote proliferation. In addition, ablation of flagellar PDEB1 and PDEB2 enhanced the parasite’s ability to colonize the hindgut of the triatomine vector, whereas PDEC-KO parasites were impaired in their establishment in the insect’s hindgut. The observed phenotypes were compartment-specific, demonstrating functional segregation between the two cAMP microdomains. Our data provide robust evidence on the presence of compartmentalized cAMP signals in *T. cruzi*, linking the role of locally synthesized cAMP pools to specific cellular responses during the parasite’s life cycle.

**Author summary:** Chagas disease is a life-threatening infectious disease caused by the protozoan parasite *Trypanosoma cruzi*, which is spread through the feces of infected kissing bugs. The parasite survives in challenging environments as it transitions between the insect vector and the mammalian host by differentiating into distinct developmental forms. cAMP is a universal second messenger that mediates specific cellular processes in the life cycle of *T. cruzi*. However, the spatial-temporal dynamics of cAMP signal remain largely unexplored in trypanosomes. We previously reported several cAMP signaling proteins in two compartments of *T. cruzi*: the contractile vacuole complex (CVC) and the flagellar tip (FT). In this study, we characterized the individual functions of these microdomains. We specifically disturbed cAMP signaling in these compartments by modulating the expression of their resident phosphodiesterases. We observed that the FT microdomain is specifically involved in parasite differentiation, host cell invasion, intracellular replication, and vector colonization, while the CVC microdomain is important for osmoregulation and parasite survival within the kissing bug. Our results unequivocally demonstrate that *T. cruzi* utilizes specific cAMP pools to address different environmental challenges. These findings highlight cAMP signaling as an essential pathway that could be further explored for the development of novel antiparasitic interventions.

## INTRODUCTION

Eukaryotic cells rely on signaling pathways to effectively detect and respond to various external stimuli through a tightly regulated process involving signal reception, amplification, and effector activation. Second messengers are central to this process, amplifying signals originating from the membrane-bound receptors. Among these, the universal second messenger 3’,5’-cyclic adenosine monophosphate (cAMP) modulates a wide variety of physiological processes in mammalian cells, including cardiac contractility [1], synaptic plasticity [2], hepatic glycogenolysis [3], adipocyte lipolysis [4], T-cell activation [5], and endocrine functions [6]. In this system, when external ligands, such as hormones or neurotransmitters, bind to G protein-coupled receptors, they initiate the activation of membrane-bound adenylate cyclases (ACs), leading to cAMP synthesis [7]. This small molecule then transduces the signal through canonical downstream effectors, including Protein Kinase A, Exchange Protein Activated by cAMP, cyclic nucleotide-gated ion channels, and Popeye domain-containing proteins [8,9]. The mechanism of cAMP signaling achieves extremely high specificity within eukaryotic cells through the activity of phosphodiesterases (PDEs), enzymes that maintain localized cAMP gradients through hydrolysis of the second messenger, while scaffold proteins enable the anchoring of signaling components to discrete subcellular compartments [10,11], thus ensuring spatial and temporal regulation of cellular responses. While this canonical pathway is conserved among eukaryotes, early divergent protozoa like trypanosomatids exhibit unique biological features that define their peculiar signaling networks [12,13].

*Trypanosoma cruzi* is the etiologic agent of Chagas disease, a vector-borne infectious disease that is considered a leading cause of disability and premature death in the Americas, for which vaccines are unavailable, and effective treatments for chronic patients are insufficient [14]. In addition, Chagas disease is spreading to non-endemic regions due to global human traveling and redistribution of vector populations [15]. Recent evidence increasingly supports that Chagas disease should be considered endemic in the United States, as *T. cruzi* infection has been detected in multiple southern states, with confirmed autochthonous human cases evidencing local transmission [16,17]. The parasite’s digenetic life cycle involves four main developmental stages, each adapted to specific niches within its insect vector (triatomine bugs) and mammalian hosts [18]. Environmental sensing is paramount for triggering developmental transitions during the parasite’s life cycle and for surviving the microenvironmental fluctuations found between and within hosts [19,20]. Different signaling pathways have been reported to modulate changes in gene expression profiles leading to morphological changes and essential physiological responses [19,21–26]. Despite recent advances in dissecting these signaling processes [27–35], the precise molecular mechanisms by which this parasite integrates diverse environmental cues to regulate differentiation and infectivity remain poorly understood.

Cyclic AMP signaling plays critical roles in trypanosomatids, reflecting the important adaptations these organisms have developed to cope with the diverse environmental challenges they encounter during their life cycles. In *Leishmania* spp., cAMP signaling has been implicated in the regulation of oxidative stress responses and the modulation of host immune evasion strategies, highlighting its importance in parasite survival [36,37]. *Trypanosoma brucei* relies on cAMP signaling to modulate social motility, the collective movement of parasite populations on semi-solid surfaces in response to extracellular cues [23,38,39], as well as mediating the parasite’s ability to colonize vector tissues [40,41], and to evade the innate immune response of the mammalian host [42]. In *Trypanosoma cruzi*, cAMP signaling has been shown to regulate two cellular processes: response to osmotic stress [26,43,44] and metacyclogenesis [27,28,45–47]. In addition, our laboratory has reported the role of cAMP signaling in other critical aspects of the *T. cruzi* life cycle, including cell adhesion, host-cell invasion, intracellular replication, and colonization of the vector’s hindgut [27,28]. Despite its biological relevance, the precise molecular mechanisms underpinning the spatiotemporal dynamics of cAMP signaling in *T. cruzi* remain unexplored.

The classical view of cAMP as a freely diffusible second messenger mediating broad cellular effects has been replaced by a model emphasizing highly organized, nanoscale signaling microdomains [48,49]. Experimental data generated using Förster resonance energy transfer (FRET)-based reporters in living cells revealed that cAMP signal specificity arises not from global cytoplasmic levels but from precisely controlled, spatially restricted gradients established by PDEs [50,51]. This principle of virtual compartmentalization, well-established in mammalian cells, is increasingly recognized in parasitic protozoa as well. Emerging evidence suggests that cAMP signals in these parasites are spatially restricted to discrete subcellular microdomains, exhibiting similarities to mammalian systems while displaying unique, parasite-specific adaptations. For instance, studies in the intestinal parasite *Giardia lamblia* have revealed compartmentalized cAMP signaling involved in encystation [52,53]. Similarly, studies in *T. brucei* have established the existence of spatially restricted cAMP signals within its flagellum [38,40,54]. Recent data from our laboratory have identified two putative cAMP microdomains in *T. cruzi*: the flagellar tip (FT) and the contractile vacuole complex (CVC) [27,28]. These two microdomains assemble cAMP signaling proteins, including specific TcAC1 and TcAC2 [27], and the multi-AC regulator cAMP response protein 3 (TcCARP3) [28]. Both studies highlight the importance of these putative microdomains for parasite survival and transmission. However, the dual localization pattern (CVC and FT) of the studied proteins (TcAC1, TcAC2, and TcCARP3) prevented the individual characterization of these microdomains. Interestingly, different PDEs, key enzymes that regulate the amplitude, duration, and degradation of cAMP, have been reported to localize at specific microdomains in *T. cruzi*, which supports our model of compartmentalized cAMP signals in this parasite [55,56].

The *T. cruzi* genome encodes five different class I PDEs: PDEA, PDEB1, PDEB2, PDEC, and PDED [57]. TcPDEB1 and TcPDEB2 localize along the flagellum [58,59], while TcPDEC localizes at the CVC [60]. Although the localization of TcPDEA and TcPDED has not been investigated, TcPDEA has been reported as a catalytically active PDE [61]. In this study, we used PDEB1, PDEB2, and PDEC as molecular markers to characterize the FT and the CVC as cAMP signaling microdomains in *T. cruzi*. Upon confirming the localization of these proteins, we generated PDEB1, PDEB2, and PDEC overexpression (OE), knockout (KO), and addback (AB) cell lines to modulate cAMP signals and investigate the individual role of these compartments. Our data indicate that cAMP generated at the CVC specifically mediates response to osmotic stress and is required for the colonization of the triatomine vector, while the FT microdomain is mainly involved in cell adhesion, metacyclogenesis, invasion of mammalian cells, and intracellular replication of amastigotes. Taken together, our results unequivocally demonstrate the importance of cAMP compartmentalized signals in the specificity of cellular responses to distinct environmental cues in the human pathogen *Trypanosoma cruzi*. Understanding the dynamics of these PDE-regulated cAMP microdomains could be key to identifying vulnerabilities in the parasite’s life cycle and potential targets for antiparasitic interventions within these signaling pathways.

## RESULTS

### Comparative topology and phylogeny of *T. cruzi* PDEs

Analysis of TcPDEs amino acid sequences revealed conserved domain architectures (Fig. 1A) that could underlie functional specialization. All five TcPDEs (TcPDEA, TcPDEB1, TcPDEB2, TcPDEC, and TcPDED) contain the characteristic C-terminal catalytic domain (PDEase, InterPro ID: IPR002073) responsible for cAMP hydrolysis. Additionally, TcPDEB1 and TcPDEB2 possess two N-terminal GAF domains (InterPro ID: IPR003018) known to bind cyclic nucleotides and regulate catalytic activity, while TcPDEC features an unusual N-terminal FYVE domain (InterPro ID: IPR000306) that is required for the localization of the protein at the CVC [43]. Structural differences, especially in the non-catalytic domains, possibly reflect the specific roles and regulatory mechanisms of each TcPDE.

**Figure 1.**
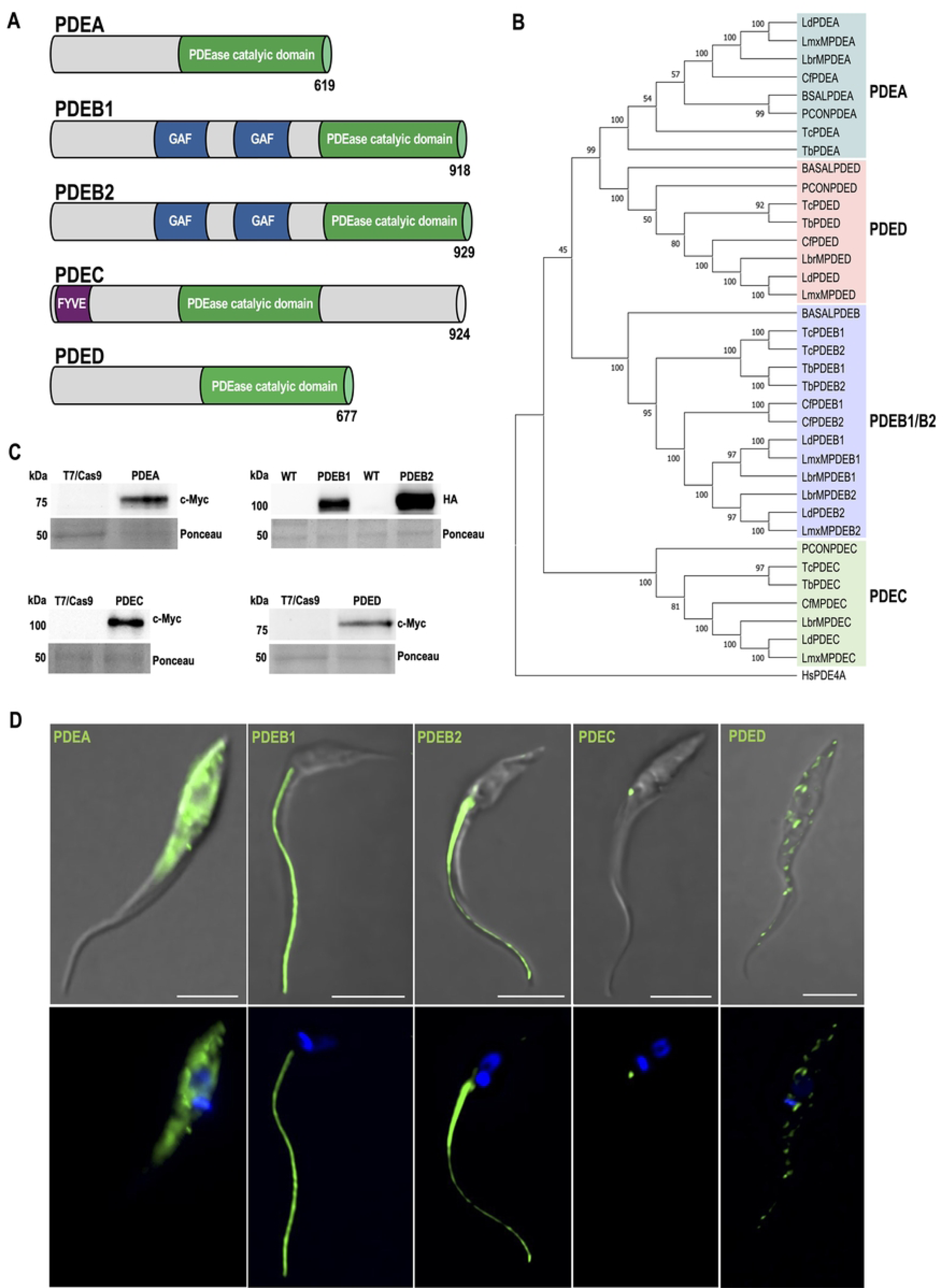
Analysis of *T. cruzi* phosphodiesterases (TcPDEs). **[A]** General topology of five TcPDEs: TcPDEA, TcPDEB1, TcPDEB2, TcPDEC, and TcPDED. TcPDEs contain a conserved C-terminal PDEase catalytic domain responsible for cAMP hydrolysis. TcPDEB1 and TcPDEB2 share 82% identity (86% similarity), contain two central GAF domains, which are predicted to bind cAMP and cGMP. TcPDEC features a unique N-terminal FYVE domain and a centrally located catalytic domain. **[B]** The phylogenetic tree was constructed using the Neighbor-Joining method, incorporating full-length PDE amino acid sequences from representative heteroxenous (*T. cruzi, T. brucei, Leishmania donovani, L. mexicana*, and *L. braziliensis*), monoxenous (*P. confusum* and *C. fasciculata*), and free-living (*B. saltans*) kinetoplastids, as well as *Homo sapiens* TcPDE4A isoform 5 (HsPDE4A). The kinetoplastid aminoacid sequences were retrieved from the publicly available genomic database, TriTrypDB, and HsPDE4A was obtained from GenBank. Gene IDs and sequence retrieval are described in *Materials and Methods*. The tree is drawn to scale, with branch lengths representing phylogenetic distances measured by the number of amino acid substitutions per site. Each sequence was labeled with an assigned gene name, and shading colors were used to distinguish four different PDE groups. **[C]** Western blot analysis of total proteins extracted from T7/Cas9 epimastigotes (control) and endogenous 3xc-Myc-tagged TcPDEA, TcPDEC, and TcPDED (76 kDa, 109 kDa, and 82 kDa, respectively) probed with anti-c-Myc antibody. Additionally, protein extracts from *T. cruzi* wild-type (WT), and HA-tagged epimastigotes overexpressing TcPDEB1 and TcPDEB2 (106 kDa and 107 kDa, respectively), were probed with anti-HA antibody. Ponceau staining served as a loading control. **[D]** Immunofluorescence assay of endogenously tagged TcPDEA, TcPDEC, and TcPDED, and TcPDEB1 and TcPDEB2 overexpressing *T. cruzi* epimastigotes. The upper panel presents the differential interference contrast (DIC) image merged with c-Myc/HA labeling (green). The lower panel displays DAPI staining (nucleus and kinetoplast, blue) merged with c-Myc/HA labeling (green). Scale bars: 5 μm.

To gain insight into the PDE families in trypanosomatids, a neighbor-joining phylogenetic tree was generated using full-length PDE amino acid sequences from representative trypanosomatids (Fig. 1B). Our dataset included heteroxenous trypanosomatids (*Trypanosoma cruzi, T. brucei, Leishmania donovani, L. mexicana*, and *L. braziliensis*), monoxenous trypanosomatids (*Paratrypanosoma confusum*and *Crithidia fasciculata*), and the free-living bodonid, *Bodo saltans*. The resulting phylogeny supports the established classification of Class I trypanosomatid PDEs into four distinct groups [57]. The analysis also revealed that all four PDE subfamilies (PDEA– PDED) are conserved in heteroxenous species (*T. cruzi, T. brucei,* and *Leishmania* spp.), while free-living *B. saltans* lacks PDEC and encodes a single PDEB isoform. In addition, *P. confusum*, the early-diverging monoxenous lineage, lacks *PDEB* orthologs, while all PDEs are present in *C. fasciculata*. The diversification of the PDE family in trypanosomatids could be an adaptation to parasitism, requiring finely tuned signaling mechanisms for survival in both mammalian hosts and vectors.

### Subcellular localization of TcPDEs supports the existence of cAMP compartments in *T. cruzi*

To investigate the possibility of spatial regulation of cAMP signaling in *T. cruzi*, we analyzed the subcellular localization of all five TcPDEs by immunofluorescence assay (IFA). For this, we generated mutant cell lines in which TcPDEA, TcPDEC, and TcPDED were endogenously tagged with a C-terminal 3xc-Myc epitope using CRISPR/Cas9 gene editing [62,63]. Due to the genomic organization in tandem and the high percentage of identity between *TcPDEB1* and *TcPDEB2* genes (87%), which prevented the CRISPR/Cas9-mediated endogenous tagging of each gene, we analyzed the individual localization of TcPDEB1 and TcPDEB2 by overexpression of C-terminal tagged versions of these proteins in *T. cruzi* epimastigotes, using pTREXn-3xHA vector [64]. Western blot analyses confirmed the expression of the five PDEs at their expected molecular weights (Fig. 1C). IFAs revealed that TcPDEC localizes to the CVC, while TcPDEB1 and TcPDEB2 localize along the flagellum (Fig. 1D), consistent with previous reports [43,58,59]. TcPDEA displayed a cytosolic distribution, while TcPDED exhibited a punctate pattern throughout the cell body (Fig. 1D). Interestingly, TcPDEA and TcPDED did not show localization patterns that overlap with those of TcPDEC or TcPDEB1 and TcPDEB2 (from now referred to as TcPDEB1/2, as both isoforms share flagellar localization). Based on their localization, we decided to use TcPDEC and TcPDEB1/2 as molecular markers to selectively modulate the content of cAMP at the cytosolic face of the CVC and FT, respectively. We named these cAMP pools CVC-cAMP and FT-cAMP. Subsequently, we confirmed the expression and localization of TcPDEC and TcPDEB1/2 in other developmental stages of *T. cruzi* (Fig. 2A-C). Our results indicate that the localization of TcPDEs was maintained across all major life cycle stages, with TcPDEC restricted to the CVC and TcPDEB1/2 to the flagellum, except for the absence of TcPDEC in metacyclic trypomastigotes (Fig. 2C). The stable localization along *T. cruzi* life cycle supports the use of TcPDEC and TcPDEB1/2 as specific markers for CVC-cAMP and FT-cAMP, respectively.

**Figure 2.**
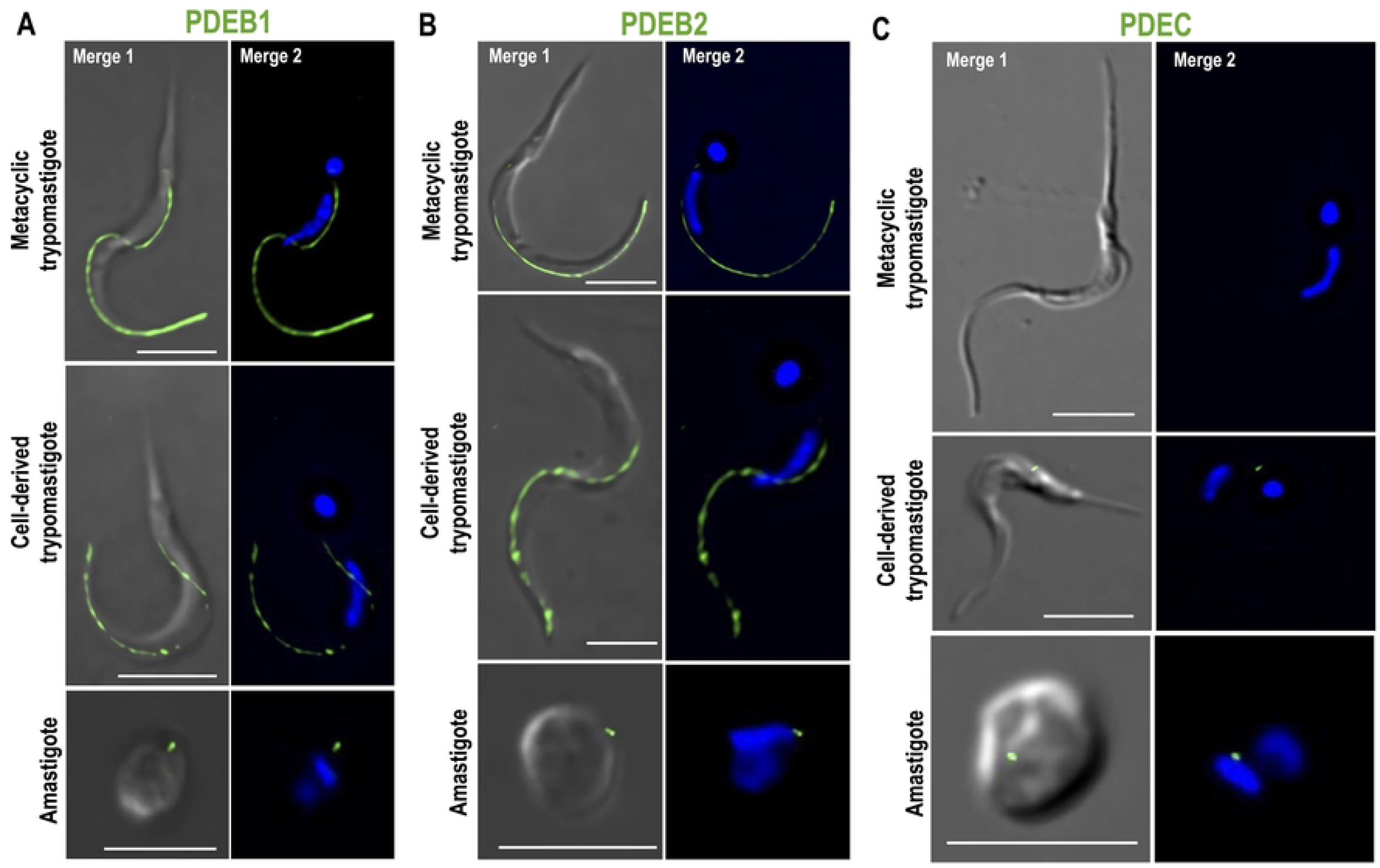
Localization of TcPDEB1, TcPDEB2, and TcPDEC in different developmental forms. Immunofluorescence assay of *T. cruzi* parasites overexpressing **[A]** TcPDEB1, **[B]** TcPDEB2, and **[C]** TcPDEC in different developmental stages as mentioned in labels at the left side of each IFA panel: metacyclic trypomastigotes, cell-derived trypomastigotes, and amastigotes. From left to right: the images show TcPDEB1-3xHA, TcPDEB2-3xHA, and TcPDEC-3xc-Myc in A, B, and C, respectively (green) merged with DIC (merge 1), or DAPI (merge 2). Scale bars: 5 μm.

### Modulation of TcPDEB1, TcPDEB2, and TcPDEC expression alters total cAMP content in *T. cruzi* epimastigotes

Following confirmation of TcPDEs localization, we used a CRISPR/Cas9-mediated gene deletion strategy [62,63] to generate a TcPDEC knockout (TcPDEC-KO) and a TcPDEB1/TcPDEB2 double knockout cell line (TcPDEB1/2-KO), to selectively modulate CVC-cAMP and FT-cAMP, respectively (Fig. 3A-D). To generate the TcPDEB1/2-KO cell line, *TcPDEB1* and *TcPDEB2* genes were simultaneously replaced as a single fragment by two resistance markers (Fig. 3C,D). Following antibiotic selection of the KO cell lines, gene ablation was confirmed by PCR (Figs. 3E,F and S1), and clonal populations were obtained by serial dilutions. The wild-type band in the parental T7/Cas9 cell line was not amplified when using external primers (Fig. 3F) due to the large (∼7.5 kb) genomic region spanning the *TcPDEB1* and *TcPDEB2* loci. However, internal primers successfully amplified the corresponding fragment from the gDNA of T7/Cas9 cells but not from *TcPDEB1/2*-KO parasites, confirming the ablation of the genes (Fig. S1E). To further confirm the ablation of *TcPDEB1/2* genes in the TcPDEB1/2-KO cell line, we used a primer set annealing on the 5’ end of the resistance markers (PAC/BSD, used to replace both alleles of *TcPDEB1/2* genes) and an external reverse primer. This primer set amplified a predicted PCR product using gDNA from TcPDEB1/2-KO parasites but not from the parental T7/Cas9 cell line (Fig. S1F,G). We then individually reintroduced *TcPDEC*, *TcPDEB1*, and *TcPDEB2* genes by cloning their open reading frames into the pTREXh-2xTy1 expression vector [28] to generate AB cell lines. Transfection of TcPDEC-KO and TcPDEB1/2-KO epimastigotes with these constructs resulted in TcPDEC-AB, TcPDEB1-AB, and TcPDEB2-AB cell lines. Expression of TcPDEC-2xTy1, TcPDEB1-2xTy1, and TcPDEB2-2xTy1 was confirmed by western blot analysis (Fig. 3G,H), while the expected localization of these tagged proteins in the AB cell lines was verified by IFA (Fig. S2A-C). In addition, we generated a TcPDEC-overexpressing cell line (TcPDEC-OE). The expression and localization of TcPDEC-OE at the CVC of epimastigotes were confirmed by western blot analysis and IFA (Fig. S2D,E).

**Figure 3.**
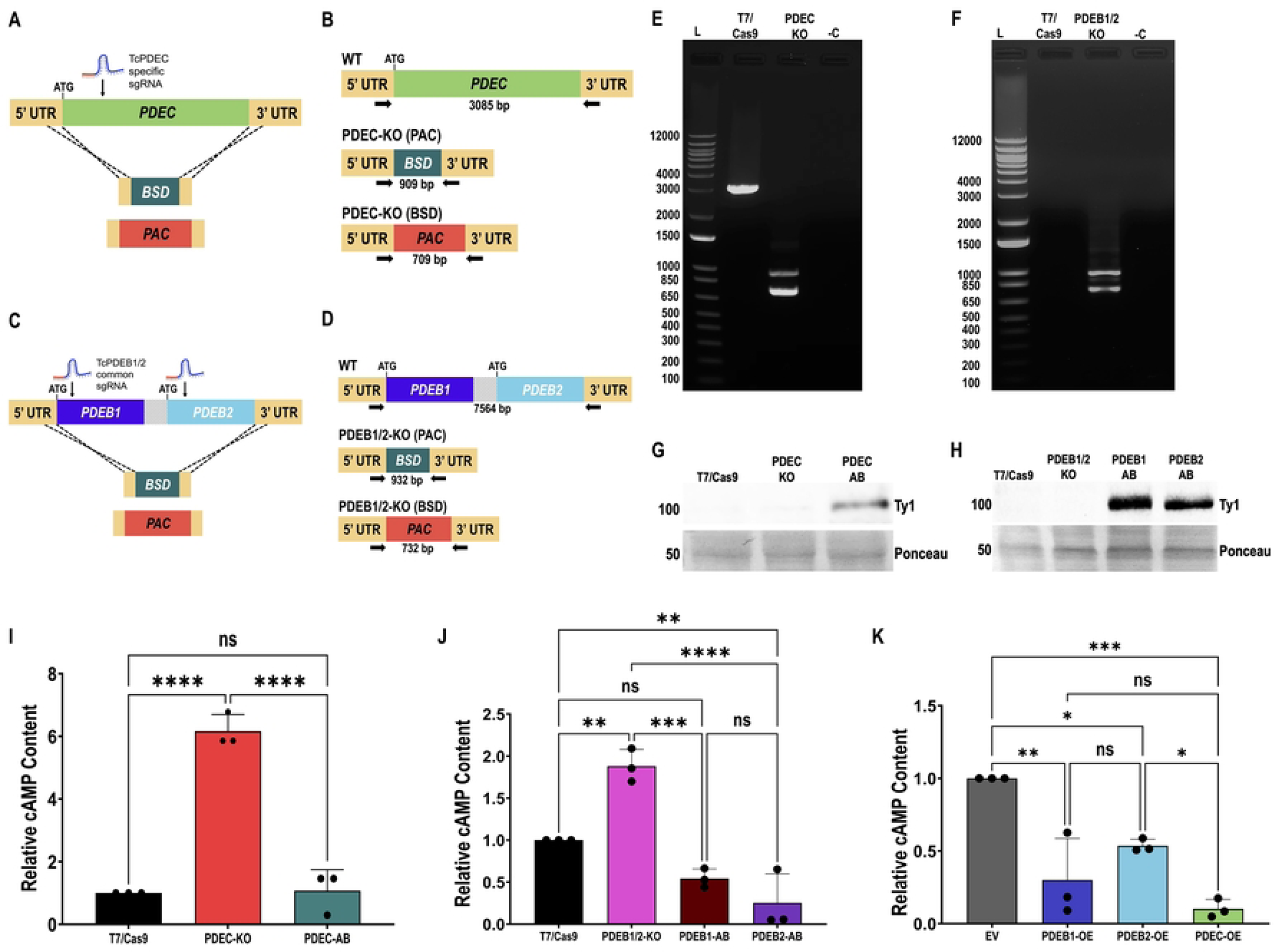
Generation of TcPDEC and TcPDEB1/2 mutant cell lines. **[A]** Schematic representation of CRISPR/Cas9-mediated strategy to generate *TcPDEC*-KO and **[B]** PCR design to verify *TcPDEC*-KO. **[C]** Schematic representation of CRISPR/Cas9-mediated strategy to generate *TcPDEB1/2* -KO and **[D]** PCR design to verify *TcPDEB1/2* -KO. **[E]** and **[F]** PCR products to verify *TcPDEC*-KO and *TcPDEB1/2*-KO resolved in 1% agarose gels. Lanen in [E]: 1 kb plus ladder (L), T7/Cas9, *TcPDEC*-KO, and negative control (-C). Lanes in [F]: 1 kb plus ladder (L), T7/Cas9, *TcPDEB1/2*-KO, and negative control (-C). **[G]** Western blot analysis confirming the expression of TcPDEC-2xTy1 (103 kDa) in the *TcPDEC*-AB cell line. T7/Cas9 and TcPDEC-KO were included as negative controls. **[H]** Western blot analysis confirming expression of TcPDEB1-2xTy1 (103 kDa) and TcPDEB2-2xTy1 (104 kDa) in the TcPDEB1-AB and TcPDEB2-AB cell lines. T7/Cas9 and TcPDEB1/2-KO were included as negative controls. Ponceau red staining was used as loading control in [G] and [H]. Intracellular cAMP content in **[I]** T7/Cas9, TcPDEC-KO and TcPDEC-AB, **[J]** T7/Cas9, TcPDEB1/2-KO, TcPDEB1-AB, and TcPDEB2-AB, and **[K]** Empty vector (EV), TcPDEB1, TcPDEB2, and TcPDEC overexpression epimastigotes. Statistical analyses were performed using one-way ANOVA with Tukey’s multiple comparisons test. Values are means ± S.D. from 3 independent experiments. *P< 0.05, **P< 0.01, ***P< 0.001.

To determine the effects of modulating TcPDEB1/2 and TcPDEC expression, we measured the cAMP content of the KO and OE cell lines alongside appropriate controls, T7/Cas9 and empty vector (EV) parasites, respectively. TcPDEC-KO parasites, lacking the CVC-localized PDE, exhibited significantly higher cAMP content than the control, and normal cAMP levels were restored in the TcPDEC-AB cell line (Fig. 3I). Similarly, TcPDEB1/2-KO parasites displayed significantly increased cAMP levels relative to control parasites, while TcPDEB1-AB and TcPDEB2-AB cell lines partially restored normal cAMP levels (Fig. 3J). Conversely, TcPDEB1-OE, TcPDEB2-OE, and TcPDEC-OE parasites showed significantly reduced cAMP content compared to the EV control (Fig. 3K). These results confirm that TcPDEB1, TcPDEB2, and TcPDEC are active cAMP-hydrolyzing enzymes. We hypothesize that TcPDEB1 and TcPDEB2 regulate the dynamics of cAMP signals generated at the FT, while TcPDEC regulates cAMP signals generated at the CVC.

### CVC-cAMP is important for epimastigote growth, while FT-cAMP regulates cell adhesion and metacyclogenesis *in vitro*

To investigate the role of compartmentalized cAMP signaling in the proliferation of *T. cruzi* epimastigotes, we monitored the *in vitro* growth of TcPDEB1/2 and TcPDEC mutant parasites. Our results indicate that TcPDEC ablation significantly impaired epimastigote growth, a defect rescued in the AB cell line (Fig. 4A,C). On the other hand, TcPDEB1/2-KO epimastigotes exhibited a mild growth defect during the exponential phase (days 2–7) compared to control parasites, and this defect was rescued in the TcPDEB1-AB and TcPDEB2-AB cell lines (Fig. 4B,D). In contrast, OE of TcPDEB1, TcPDEB2, or TcPDEC did not significantly impact the growth of epimastigotes (Fig. S3A). These results suggest that increased levels and free diffusion of CVC-cAMP impair epimastigote proliferation, highlighting the importance of this putative cAMP compartment for cell physiology.

**Figure 4.**
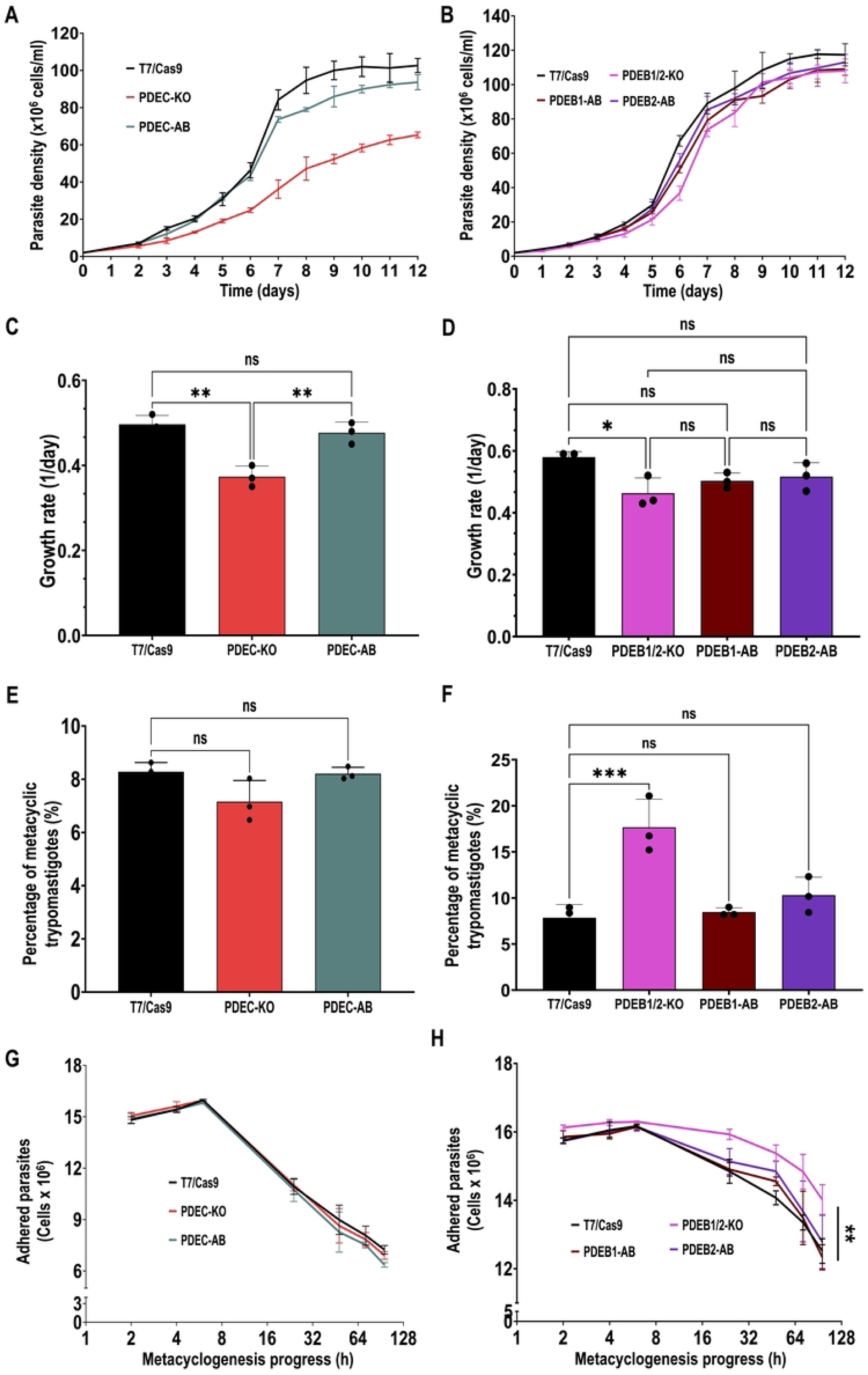
Growth, metacyclogenesis, and adhesion phenotype. Growth curve of **[A]** T7/Cas9, TcPDEC-KO, TcPDEC-AB epimastigotes and **[B]** T7/Cas9, TcPDEB1/2-KO, TcPDEB1-AB, TcPDEB2-AB epimastigotes in LIT medium. **[C]** and **[D]** shows the growth rate of cell lines in [A] and [B] during the exponential phase (days 2-7). **[E]** and **[F]** shows the in vitro metacyclogenesis of cell lines in [A] and [B]. Values are means ± S.D., n=3. Statistical analysis was performed using one-way ANOVA with Dunnett’s multiple comparisons. **[G]** and **[H]** shows adhesion assays using cell lines in [A] and [B]. Values are means ± S.D., n=3. Statistical analysis was performed using two-way ANOVA with Dunnett’s multiple comparisons. *P< 0.05, **P< 0.01, ***P< 0.001, ****P< 0.0001.

Next, we evaluated the role of compartmentalized cAMP signaling in *T. cruzi* metacyclogenesis by assessing the ability of mutant cell lines to differentiate *in vitro* as described in *Materials and Methods*. Our results indicate that TcPDEB1/2-KO parasites exhibited a significantly higher percentage of metacyclic trypomastigotes than control cells, a defect that was fully restored in the TcPDEB1-AB and TcPDEB2-AB cell lines (Fig. 4F). Conversely, TcPDEC-KO parasites did not show a defect in metacyclogenesis (Fig. 4E), suggesting that only FT-cAMP signals modulate this process.

Given that TcPDEB1/2-KO parasites exhibited enhanced metacyclogenesis, we examined their ability to adhere to culture flask surfaces, as adhesion was previously reported as a key process for metacyclogenesis [65]. Indeed, in agreement with their enhanced differentiation, TcPDEB1/2-KO parasites showed significantly increased adhesion, a phenotype that was fully rescued in the AB cell lines (Fig. 4H). Conversely, TcPDEC ablation did not impact adhesion (Fig. 4G), supporting the hypothesis that only FT-cAMP signals (and not CVC-cAMP) modulate this process. Interestingly, enhanced adhesion and metacyclogenesis were evident only when the FT-cAMP pool was increased (PDEB1/2-KO mutant), while parasites with reduced cAMP levels (OE cell lines) did not exhibit significant changes in these processes (Fig. S3B,C), indicating that FT-cAMP is a positive modulator of metacyclogenesis. Together, these results confirm that adhesion is a prerequisite for metacyclogenesis and that FT-cAMP signals, but not CVC-cAMP, mediate both processes in *T. cruzi*.

### CVC-cAMP specifically mediates the response to hypoosmotic stress

To determine if CVC-cAMP signaling exclusively mediates the response to hypoosmotic stress, we performed regulatory volume decrease (RVD) assays on the mutant cell lines as described in *Materials and Methods*. We evaluated RVD capacity based on maximum cell volume change upon hypoosmotic stress (from 282 to 115 mOsmol/L) and final volume recovery after 12 minutes. TcPDEC-KO parasites showed reduced cell swelling and more efficient initial volume recovery compared to the T7/Cas9 control, while the phenotype was fully restored in TcPDEC-AB parasites, suggesting that enhanced RVD resulted from increased CVC-cAMP levels (Fig. 5A-C). These results indicate a positive correlation between high CVC-cAMP and enhanced osmoregulatory capacity. Conversely, TcPDEB1/2-KO parasites showed no significant differences in RVD compared to control and AB cell lines, suggesting that FT-cAMP signals are not involved in this process (Fig. 5D-F). To further explore the specificity of cAMP compartments, we performed RVD assays with PDE-OE parasites. Our results indicate that TcPDEC-OE epimastigotes exhibited increased swelling and failed to recover their volume upon hypoosmotic stress (Fig. 5G-I). In contrast, TcPDEB1-OE and TcPDEB2-OE parasites did not show significant differences in their osmoregulatory capacity compared to the EV control, further confirming that CVC-cAMP signals, but not FT-cAMP, specifically regulate the ability of *T. cruzi* epimastigotes to recover their cell volume under hypoosmotic conditions.

**Figure 5.**
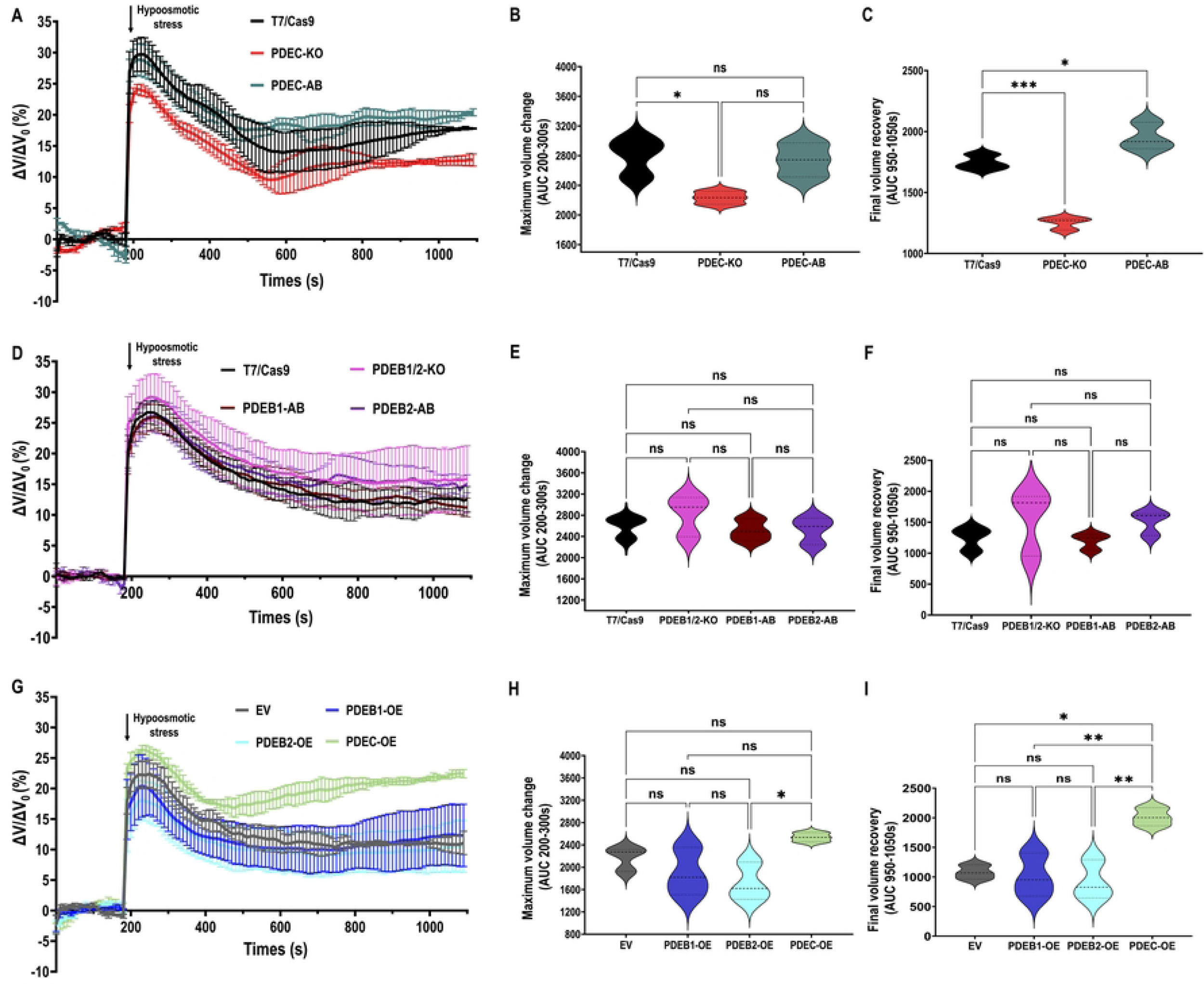
Regulatory Volume Decrease (RVD) after hypoosmotic stress. **[A]** The light scattering pattern of *T. cruzi* epimastigotes suspended in isotonic buffer was recorded for 120 s and diluted to a final osmolarity of 115 mOsm/L under constant ionic conditions. Relative changes in cell volume were monitored by measuring absorbance at 550 nm over time in T7/Cas9, TcPDEC-KO, and TcPDEC-AB epimastigotes. The absorbance values were normalized to the initial volume under isosmotic conditions and expressed as the percentage of volume change. **[B]** Analysis of the maximum volume change under hypoosmotic conditions. The area under the curve (AUC) in [A] was calculated between 200 and 300 seconds for the cell lines. **[C]** Final volume recovery was calculated as the AUC in [A] between 950 and 1050 seconds. **[D]**, **[E]**, and **[F]** The same experiments as in [A], [B], and [C] with T7/Cas9, TcPDEB1/2-KO, TcPDEB1-AB, and TcPDEB2-AB epimastigotes. **[G]**, **[H]**, and **[I]** The same experiments as in [A], [B], and [C] with Empty Vector, TcPDEB1-OE, TcPDEB2-OE, and TcPDEC-OE cell lines. Statistical analysis was performed using one-way ANOVA with Tukey’s multiple comparison test. Values are mean ± SD; n = 3. *P< 0.05, **P< 0.01, ***P< 0.001, ns = not significant differences.

### FT-cAMP signals are necessary for host cell invasion and intracellular replication

To determine the role of compartmentalized cAMP signaling in the invasion of mammalian cells, we assessed the ability of TcPDE mutant cell-derived trypomastigotes to infect human foreskin fibroblasts (hFFs) as described in *Materials and Methods*. The percentage of infected cells was analyzed by fluorescence microscopy of 4′,6-diamidino-2-phenylindole (DAPI)-stained hFFs 24 h post-infection. Our results indicate that TcPDEB1/2-KO parasites exhibit a defect in the ability to infect hFFs, as compared to T7/Cas9 parasites. This phenotype was rescued in the TcPDEB2-AB cell line (Fig. 6A), which we chose as rescued control because it showed a lower cAMP content than TcPDEB1-AB parasites (Fig. 3J). We next quantified the number of amastigotes per infected host cell at 72 h post-infection and found that TcPDEB1/2-KO parasites also exhibited a significant intracellular replication defect that was restored in TcPDEB2-AB parasites (Figs. 6B and S4). Interestingly, TcPDEB1-OE and TcPDEB2-OE parasites also displayed an impaired ability to invade hFFs but showed no defects in intracellular replication (Fig. 6C,D). These findings indicate that any alteration in basal FT-cAMP levels impairs the parasite’s ability to invade mammalian cells. In contrast, modulating CVC-cAMP levels (TcPDEC-KO and TcPDEC-OE) produced no significant defects in either invasion or intracellular replication (Fig. 6E-H), increasing the body of evidence that supports the specificity of the FT-cAMP microdomain in regulating *T. cruzi* infectivity.

**Figure 6.**
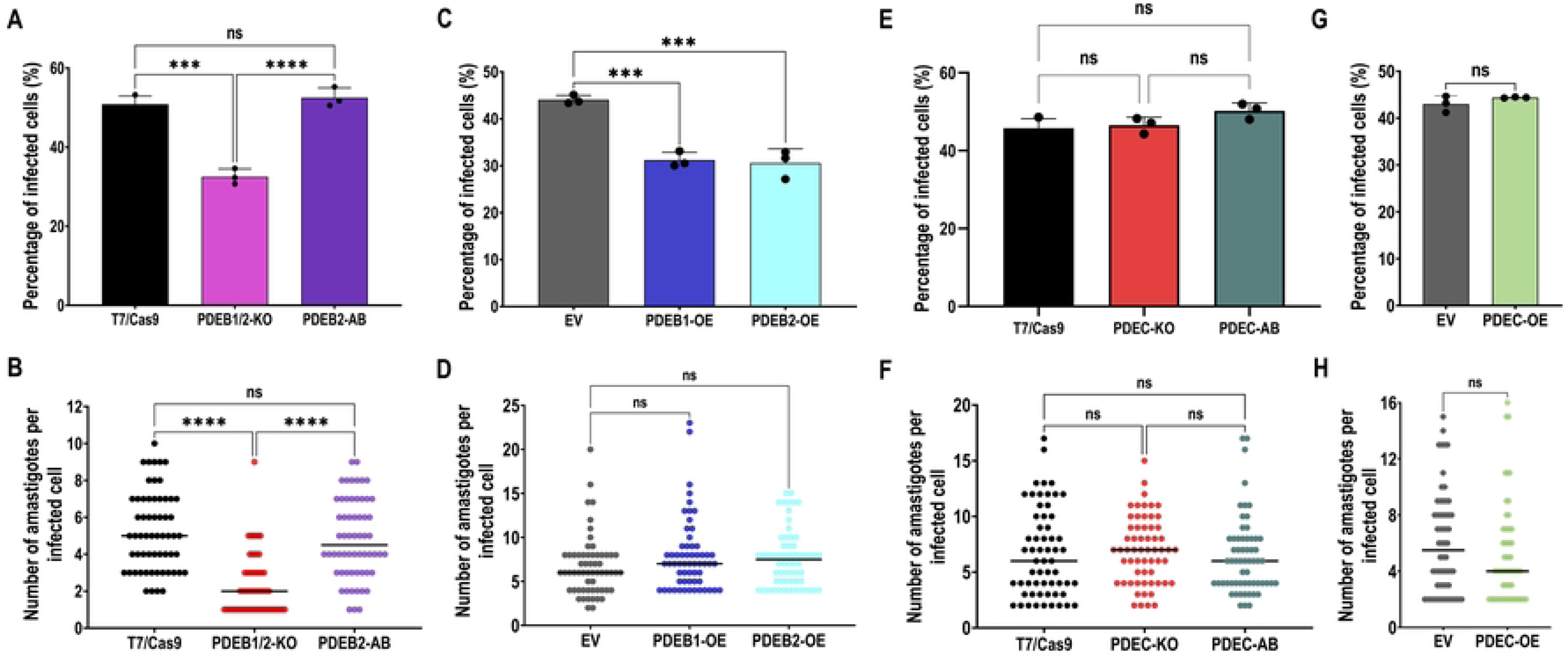
Mammalian cell invasion and intracellular replication. The percentage of infected hFFs 24h post-infection was evaluated in **[A]** T7/Cas9, TcPDEB1/2-KO, and TcPDEB2-AB, **[C]** Empty vector, TcPDEB1-OE, and TcPDEB2-OE, **[E]** T7/Cas9, TcPDEC-KO, and TcPDEC-AB, **[G]** Empty vector and TcPDEC-OE trypomastigotes. Values are means ± S.D., n=3. Statistical analysis was performed using one-way ANOVA with Tukey’s multiple comparisons tests in [A], [C], and [E] and Student’s t-test in [G]. The number of intracellular amastigotes per infected host cell 72 h post-infection was evaluated in **[B]** T7/Cas9, TcPDEB1/2-KO, and TcPDEB2-AB, **[D]** Empty vector, TcPDEB1-OE, and TcPDEB2-OE, **[F]** T7/Cas9, TcPDEC-KO, and TcPDEC-AB, and **[H]** Empty vector and TcPDEC-OE trypomastigotes. The black line indicates the median value in each cell line, n=60. Statistical analysis was performed using the Kruskal-Wallis test with Dunn’s multiple comparisons in [B], [D], [F], and [H]. *P< 0.05, **P< 0.01, ***P< 0.001, ****P< 0.0001.

### CVC-cAMP and FT-cAMP differentially modulate the parasite’s ability to colonize the hindgut of the triatomine vector

To determine how compartmentalized cAMP signaling influences *T. cruzi* transmission, we infected the triatomine bug *Rhodnius prolixus* with TcPDE knockout and control cell lines, as described in *Materials and Methods*. Upon dissection, the infected bugs were distinguished from non-infected ones by microscopic analysis of hindguts to visualize epimastigotes and metacyclic trypomastigotes in this portion of the vector’s digestive tract. Our results revealed contrasting colonization phenotypes. The percentage of kissing bugs containing *T. cruzi* parasites in their hindguts was significantly higher when infected with TcPDEB1/2-KO parasites, suggesting an enhanced vector colonization phenotype (Fig. 7A). This result is consistent with their increased adhesion and metacyclogenesis phenotypes *in vitro*, with the AB lines restoring the normal phenotype. In contrast, TcPDEC-KO parasites displayed a significantly reduced kissing bug colonization compared to control and AB cell lines (Fig. 7B). We found these results very intriguing, as these parasites showed normal metacyclogenesis and adhesion phenotypes (Fig. 4E-G). Furthermore, despite being more tolerant to hypoosmotic stress in RVD assays, TcPDEC-KO parasites showed a reduced colonization of the vector’s hindgut. A possible explanation is that these parasites do not tolerate the hyperosmotic conditions in the vector’s hindgut, which increases from 300-400 to 700-900 mOsm/kg two to three days after a bloodmeal [66]. To test this hypothesis, we performed a regulatory volume increase assay under hyperosmotic conditions by exposing *T. cruzi* epimastigotes to a hypertonic buffer to increase the osmolarity from 282 to 860 mOsmol/L, as described in *Materials and Methods*. Interestingly, we observed that TcPDEC-KO parasites experienced a significantly higher shrinkage than control and AB cell lines upon hyperosmotic stress (Fig. 7C, D), being unable to recover their normal volume even after 15 min. This impaired ability to regulate cell volume in a hyperosmotic environment could explain their failure to survive in the harsh environment of the triatomine hindgut. Taken together, our vector colonization results unveiled specific roles for CVC and FT cAMP compartments on parasite transmission, which could be relevant for the development of new strategies for antiparasitic interventions.

**Figure 7.**
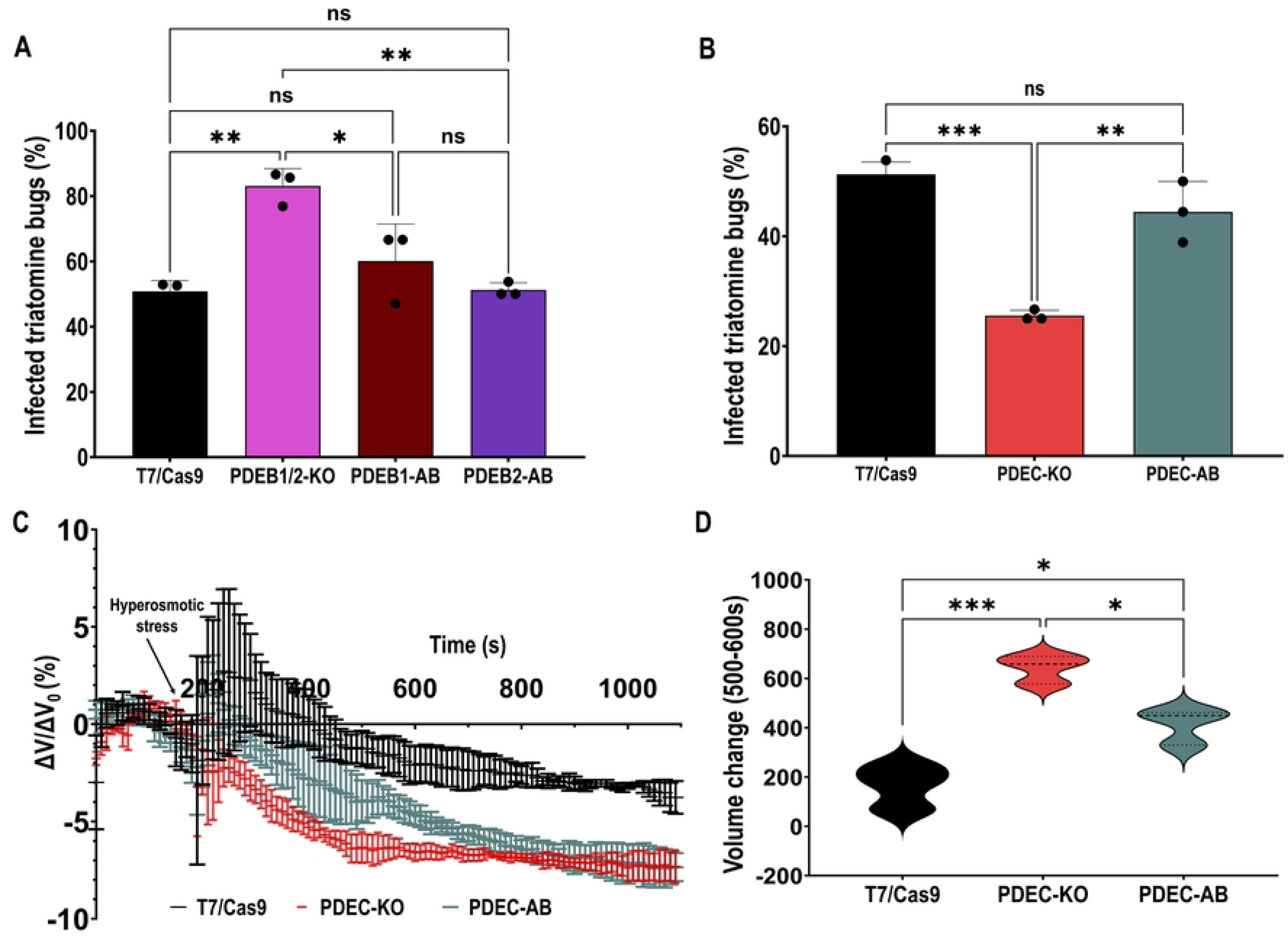
Kissing bug colonization by TcPDE knockout parasites. Percentage of infected triatomine bugs. Kissing bugs were infected by feeding blood containing **[A]** T7/Cas9, TcPDEB1/2-KO, TcPDEB1-AB, and TcPDEB2-AB epimastigotes, and **[B]** T7/Cas9, TcPDEC-KO, and TcPDEC-AB epimastigotes. The insects’ hindguts were dissected 30 days post-feeding and examined under the microscope for the presence of parasites. Values are mean ± S.D., n=3 independent experiments with 9-15 insects per group infected with each cell line. Statistical analysis was performed using one-way ANOVA with Tukey’s multiple comparisons. *P< 0.05, **P< 0.01, ***P< 0.001. **[C]** Regulatory volume responses upon hyperosmotic stress. After 3 min under isosmotic conditions, cells were exposed to hyperosmotic stress at a final osmolarity of 860 mOsm/L. **[D]** Volume changes were calculated as explained in [Figure 6]. The area under the curve (AUC) in [C] was calculated between 500 and 600 seconds for the cell lines. Statistical analysis was performed using one-way ANOVA with Tukey’s multiple comparisons. Values are mean ± SD; n = 3. *P< 0.05, **P< 0.01, ***P< 0.001.

## Discussion

This study provides the first evidence for functionally specialized cAMP signaling microdomains in *T. cruzi*. Building upon our previous identification of two putative cAMP microdomains in this parasite [27,28], we overcame the interpretative challenges posed by the dual localization of TcAC1 and TcCARP3 by capitalizing on the specific localizations of TcPDEC at the CVC and TcPDEB1/TcPDEB2 along the flagellum. Our data indicate that CVC-cAMP specifically mediates response to osmotic stress and is required for the colonization of the triatomine vector, while FT-cAMP is involved in cell adhesion, metacyclogenesis, invasion of mammalian cells, and intracellular replication of amastigotes. Overall, our results provide robust evidence on the role of cAMP compartmentalized signals in the specificity of cellular responses to different environmental cues in the digenetic parasite *T. cruzi*.

PDE localizations were consistent across the main developmental stages of the parasite, except for TcPDEC, which remained undetectable in metacyclic trypomastigotes. This observation aligns with our previous report of CVC localization for TcAC1 and TcCARP3 [28]. This result could be explained by the smaller size of the CVC in metacyclic trypomastigotes than in other developmental stages [67], possibly placing these CVC-resident proteins below the sensitivity threshold of epifluorescence microscopy. In addition, TcPDEC expression may be downregulated or absent in this developmental stage, since metacyclic trypomastigotes develop in a naturally hyperosmotic environment, the hindgut of the triatomine bug, where the RVD machinery of the CVC is not needed for cell survival [68]. On the other hand, TcPDEA and TcPDED showed specific subcellular localizations (cytosolic and undetermined punctate pattern, respectively), different from that of PDEC and PDEBs (CVC and flagellum, respectively), suggesting that each PDE may regulate a spatially confined cAMP pool within the cell. The experimental approach of using TcPDEC and TcPDEB1/2 as molecular markers for the CVC and the FT microdomains allowed us to selectively modulate cAMP levels within these two previously identified cAMP compartments. Modulating the gene expression of *TcPDEB1/2* and *TcPDEC* by KO and OE approaches allowed us to dissect individual contributions of CVC-cAMP and FT-CAMP signaling and resolve ambiguities regarding their functional specificity. These findings significantly advance our understanding of how *T. cruzi* senses environmental cues and responds to stress conditions by utilizing locally synthesized cAMP signals.

The cAMP content data provided additional evidence that TcPDEC, TcPDEB1, and TcPDEB2 are catalytically active PDEs that modulate intracellular cAMP levels in *T. cruzi*, as previously reported [43,58–60,69]. The elevated cAMP content observed in the TcPDEC-KO and TcPDEB1/2-KO mutants confirmed the hydrolytic nature of these enzymes, while the partial or even total restoration of basal cAMP levels in the corresponding AB cell lines supports the specificity of each of these TcPDEs in maintaining cAMP homeostasis. Likewise, OE of TcPDEC, TcPDEB1, and TcPDEB2 led to significant reductions in the total content of cAMP, further supporting their role in cAMP hydrolysis. These results encouraged us to use TcPDEC and TcPDEB1/2 as molecular markers to selectively modulate CVC-cAMP and FT-cAMP levels, respectively.

Our data revealed that CVC-cAMP, modulated by TcPDEC, specifically mediates osmoregulation via RVD under hypoosmotic conditions. TcPDEC-KO parasites displayed enhanced tolerance to hypoosmotic stress, confirming the existence of a cAMP-dependent mechanism of volume recovery involving the CVC compartment. These findings support a model of cAMP-mediated RVD in *T. cruzi* [44], in which cAMP generated at the cytosolic face of the CVC triggers the tubulin-mediated fusion of acidocalcisomes to the central vacuole of the CVC, with the consequent translocation of the aquaporin TcAQP1 to the CVC membrane, and simultaneous release of the acidocalcisome content into the central vacuole. Subsequently, TcAQP1 allows water uptake by the central vacuole of the CVC. Pulsatile contractions of the central vacuole facilitate fluid discharge through the flagellar pocket via adhesion plaque remodeling [70], leading to cell volume recovery upon hypoosmotic stress. The increased tolerance to hypoosmotic stress observed in TcPDEC-KO parasites is consistent with previous results in *T. cruzi* epimastigotes showing that chemical inhibition of TcPDEC enhances RVD under hypoosmotic stress [43,60]. These results are also consistent with the osmotolerance phenotype observed in TcAC1-overexpressing parasites [27]. The strong effect of TcPDEC ablation on the osmoregulatory capacity of epimastigotes was further illustrated by the RVD defect observed in TcPDEC-OE parasites. These results are consistent with previous reports where TcPDEC-overexpressing cells showed a defect in volume recovery, becoming insensitive to PDE inhibitors [43,60]. More importantly, the RVD phenotypes exhibited by these PDC mutants were not observed in PDEB1/2-KO, AB, and overexpressing parasites, suggesting that this cellular process is specifically regulated by CVC-cAMP. Together, these results strongly support the existence of spatially regulated CVC-cAMP signals that are critical for adaptive responses to the fluctuating osmolarities encountered during parasite transitions between and within hosts. Interestingly, elevated CVC-cAMP in TcPDEC-KO parasites significantly impaired the proliferation of epimastigotes, revealing additional cAMP-mediated metabolic functions that should be further investigated. The growth defect we observed in TcPDEC-KO parasites is in agreement with previous data from our group, showing that TcCARP3-KO epimastigotes also display impaired growth [28]. Here, we provide evidence on the contribution of CVC-cAMP-altered signals to this growth defect. Importantly, the specificity of these phenotypes confirms functional segregation between CVC and FT as cAMP microdomains.

The flagellar distal domain (flagellar tip) has been proposed as a cAMP signaling microdomain in *T. cruzi* [27,28]. Our functional characterization of the FT-cAMP compartment, regulated by TcPDEB1 and TcPDEB2, validates it as a separate signaling hub regulating cell adhesion in epimastigotes, differentiation to metacyclic trypomastigotes, host cell invasion by cell-derived trypomastigotes, and intracellular replication of amastigotes. Parasites lacking TcPDEB1 and TcPDEB2 showed increased adhesion and enhanced metacyclogenesis *in vitro*, suggesting that FT-cAMP signals are involved in developmental processes critical for parasite transmission. The importance of flagellar PDEs in regulating environmental responses has been widely reported in trypanosomes [38,40,54]. In *T. brucei*, simultaneous RNAi knockdown of TbPDEB1/2 in bloodstream forms disrupts normal cell division, leading to parasite death [71], which differs to our findings in *T. cruzi,* where a mild growth defect was observed in the TcPDEB1/2-KO cell line. These contrasting outcomes suggest parasite-specific differences in the role of flagellar PDEs between *T. cruzi* and *T. brucei*. Furthermore, in *T. brucei*, flagellar PDEB1 is essential for sensing the environment and navigating through the tsetse fly vector, as ablation of this protein results in a pronounced defect in social motility, compromising the parasite’s capacity to migrate to the salivary glands [40]. Interestingly, while PDEB1/2 ablation promotes adhesion and differentiation in *T. cruzi*, the opposite outcome has been observed in other kinetoplastids. In *C. fasciculata*, pharmacological inhibition of PDEs, which increases the intracellular levels of cAMP, completely blocks the parasite’s ability to attach to a substrate [72]. Similarly, in *Trypanosoma congolense*, parasite treatment with a PDE inhibitor promotes the detachment of parasites from surfaces [73]. These observations suggest that whereas the flagellum is a conserved signaling hub across kinetoplastids, the downstream effects of cell signaling have been adapted to suit the unique biological features of each species, leading to diverse attachment mechanisms in trypanosomatids [65,74,75]. Despite these differences, the role of FT-cAMP in parasite attachment and differentiation connects several foundational concepts: that physical contact of the tip of the flagellum to a substrate is a key trigger for metacyclogenesis [65,76], the role of cAMP as a molecular inducer of differentiation [46,47]; and that environmental cues like nutrient deprivation initiate metacyclogenesis through still unknown mechanisms that lead to activation of ACs in *T. cruzi* [45].

We have previously reported the role of cAMP in essential processes for parasite development in the mammalian host [27,28]. In the present study, we observed that tightly regulated levels of cAMP synthesized in the FT-cAMP are essential for parasite infectivity, as TcPDEB1/2 ablation or OE impaired host cell invasion and intracellular replication. The fact that FT-cAMP specifically governs parasite infectivity explains our previous observations with dually localized proteins TcAC1 [27] and TcCARP3 [28]. In those studies, modulating the expression of TcAC1 and TcCARP3 altered cAMP levels simultaneously in both the FT and the CVC microdomains, which made the interpretation of microdomain-specific roles more difficult. Conversely, we now use TcPDEB1 and TcPDEB2, as exclusive regulators of FT-cAMP, which allowed us to recognize host cell invasion and intracellular replication as specific processes orchestrated by this microdomain. In summary, our data suggests that the flagellar distal domain is a structurally and functionally defined signaling compartment that is specifically involved in cell adhesion, differentiation, host cell invasion, and intracellular replication in *T. cruzi*. Importantly, FT-cAMP perturbation minimally affected epimastigote proliferation and had no detectable impact on osmoregulation, further emphasizing clear functional segregation from the CVC-cAMP microdomain.

The role of cAMP in vectorial transmission has been previously reported in trypanosomes, finding this signaling pathway to be essential for colonization of the insect vector by trypanosomes with either salivarian transmission (*T. brucei*) and stercorarian transmission (*T. cruzi*) [28,40,54,77]. Here, we explored how compartmentalized cAMP signaling impacts parasite transmission via its insect vector, *Rhodnius prolixus*. TcPDEB1/2-KO parasites showed significantly enhanced ability to colonize the insect hindgut, a phenotype that is concomitant with the increased cAMP content, parasite adhesion, and metacyclogenesis displayed by this mutant, where FT-cAMP is expected to be elevated. Conversely, despite normal adhesion and differentiation capacities, TcPDEC-KO parasites exhibited an impaired ability to colonize the hindgut of kissing bugs. Although these parasites show an enhanced response to hypoosmotic conditions, they display increased shrinkage under hyperosmotic stress compared to control cells, which is a critical physiological challenge this parasite encounters in the vector’s hindgut [78,79]. Excessive shrinkage of TcPDEC-KO parasites under hyperosmotic conditions suggests impaired regulatory volume increase essential for survival in the hindgut environment, a phenotype partially rescued in TcPDEC-AB parasites. These findings align with previous reports [22,26,44,68,79–82], showing that the CVC and acidocalcisomes act in coordination to sense and respond to osmotic fluctuations in *T. cruzi*, particularly in the hindgut, where osmolarities exceed 750 mOsm/kg [66], and upon infecting the mammalian host, where osmolarities decrease to physiological levels (300 mOsm/kg [19]. Several studies have reported that hyperosmotic stress triggers rapid cell shrinkage and CVC enlargement, followed by compensatory responses, such as polyphosphate synthesis and amino acid accumulation [22,79]. Furthermore, the discovery of TcMscS, a lipid-activated mechanosensitive ion channel in the central vacuole of the CVC [35,83], supports the hypothesis that CVC-localized sensors modulate adaptive responses to membrane alterations induced by osmotic stress. The stercorarian transmission mode of *T. cruzi*, which depends on successful colonization of the insect hindgut, makes osmotic resilience uniquely critical. In contrast, the salivarian trypanosome *T. brucei* is not exposed to such selective pressure, which could have driven the loss of the CVC in evolution [22,68]. Here, we are reporting for the first time a triatomine colonization defect associated with impaired osmoregulation in *T. cruzi*.

Our findings on compartmentalized cAMP signals in *T. cruzi* agree with a broader paradigm shift in eukaryotic cAMP signaling, moving away from classical models of cAMP as a diffusible second messenger towards recognizing cAMP as spatially confined within virtual microdomains. Such a concept of localized signaling domains has been established largely due to seminal work in mammalian systems [48,50,51]. For example, FRET-based biosensors in cardiomyocytes have been used to demonstrate the existence of discrete cAMP nanodomains [51], where local signals shaped by the activity of PDEs are more important for the regulation of specific cell functions than the global cytosolic pool of cAMP. This work established the concept that the spatial restriction of cAMP, controlled by the precise localization of cAMP signaling proteins (ACs, PDEs, and effectors/modulators), is a key mechanism for determining signal specificity. Our results support this model in a protozoan parasite, where the exclusive localization of specific PDEs to either the CVC or the FT creates functionally isolated signaling hubs. Comparable compartmentalization strategies are apparent in other protozoa, including *Giardia lamblia* [52,53] and *T. brucei* [38,40]. However, our study identifies unique functional specializations linking FT-cAMP signals to parasite differentiation and infectivity, and CVC-cAMP signals to osmoregulatory capacity, reflecting the specific evolutionary pressures shaping the biology of *T. cruzi*.

Our phylogenetic analysis of Class I PDEs across kinetoplastids suggests that the compartmentalization of cAMP signals in trypanosomatids could have emerged in response to parasitic lifestyles. While all four PDE subfamilies (PDEA–PDED) are conserved in parasitic trypanosomatids (*T. cruzi*, *T. brucei*, *C. fasciculata*, and *Leishmania* spp.) [21,24], PDEC is absent in the free-living bonodid *B. saltans* [84], and PDEB was not retained in the early-diverging monoxenous kinetoplastid *P. confusum* [85]. This phylogenetic distribution suggests that PDE family expansion and specialization is an adaptations to parasitism, reflecting selective pressures imposed by host environments. The presence of both PDEB isoforms across all parasitic trypanosomatids except in the early-diverging *P. confusum*, highlights the essential role of FT-cAMP microdomain in regulating critical transmission processes, which is supported by our results of TcPDEB1/2-mediated control of adhesion, metacyclogenesis, host-cell invasion, and intracellular replication. Similarly, the emergence of PDEC in parasitic lineages, in contrast to its absence in free-living bodonids, correlates with its specialized role in CVC-mediated osmoregulation, which is crucial for surviving the dramatic osmotic fluctuations encountered within insect and mammalian hosts [44,86]. Together, these findings support the idea that compartmentalized cAMP signaling in *T. cruzi* is not a conserved feature, but rather a sophisticated, evolutionarily refined system where PDE isoform localization and functional diversification precisely meet the signaling demands of a complex parasitic existence. By integrating phylogenetic analysis with cellular physiology evidence, our work supports the concept of compartmentalized cAMP signaling as an adaptation to parasitism in trypanosomes.

The specific phenotypes observed in our PDE mutants strongly support the existence of compartmentalized cAMP signaling. However, our measurements relied on total cAMP content rather than on compartment-specific cAMP levels. Analysis of this dynamics will require the development of a genetically encoded, fluorescence or FRET-based cAMP biosensor for *T. cruzi*, similar to those developed in mammalian cells [48,51], *T. brucei* [39], and *Giardia* [52]. Alternatively, characterization of TcPDEA and TcPDED could unveil their roles as putative regulators of compartmentalized cAMP signaling in *T. cruzi*.

Our results contribute to a body of evidence [27,28,38–41,52,55,56,87] supporting a model of environmental sensing where external cues such as osmotic stress, nutrient deprivation, pH shifts, temperature changes, or cell contact may alter membrane fluidity and lipid raft composition, thereby triggering the activation of specific ACs in *T. cruzi*. Given the dual localization of TcAC1 and TcAC2 to both the FT and the CVC, we hypothesize that these enzymes function as non-canonical environmental sensors capable of selectively activating distinct cAMP microdomains in response to specific stimuli. This membrane-dependent activation could initiate spatially restricted cAMP signaling cascades that mediate specific cellular responses. Figure 8 illustrates our current model of functional segregation of cAMP compartments in *T. cruzi*, which aligns with emerging evidence from other protozoan systems where membrane dynamics replaces classical G protein-coupled receptor-mediated cAMP signaling [52,53,87] and highlights the peculiar features of kinetoplastid parasites in evolving non-conserved signal transduction mechanisms to complete their complex life cycles. Further research is necessary to identify the TcAC activation mechanism in each microdomain and the downstream effectors that transduce their specific signals, which will be instrumental in advancing our understanding of cAMP compartmentalization and its role in environmental sensing and parasite survival.

**Figure 8.**
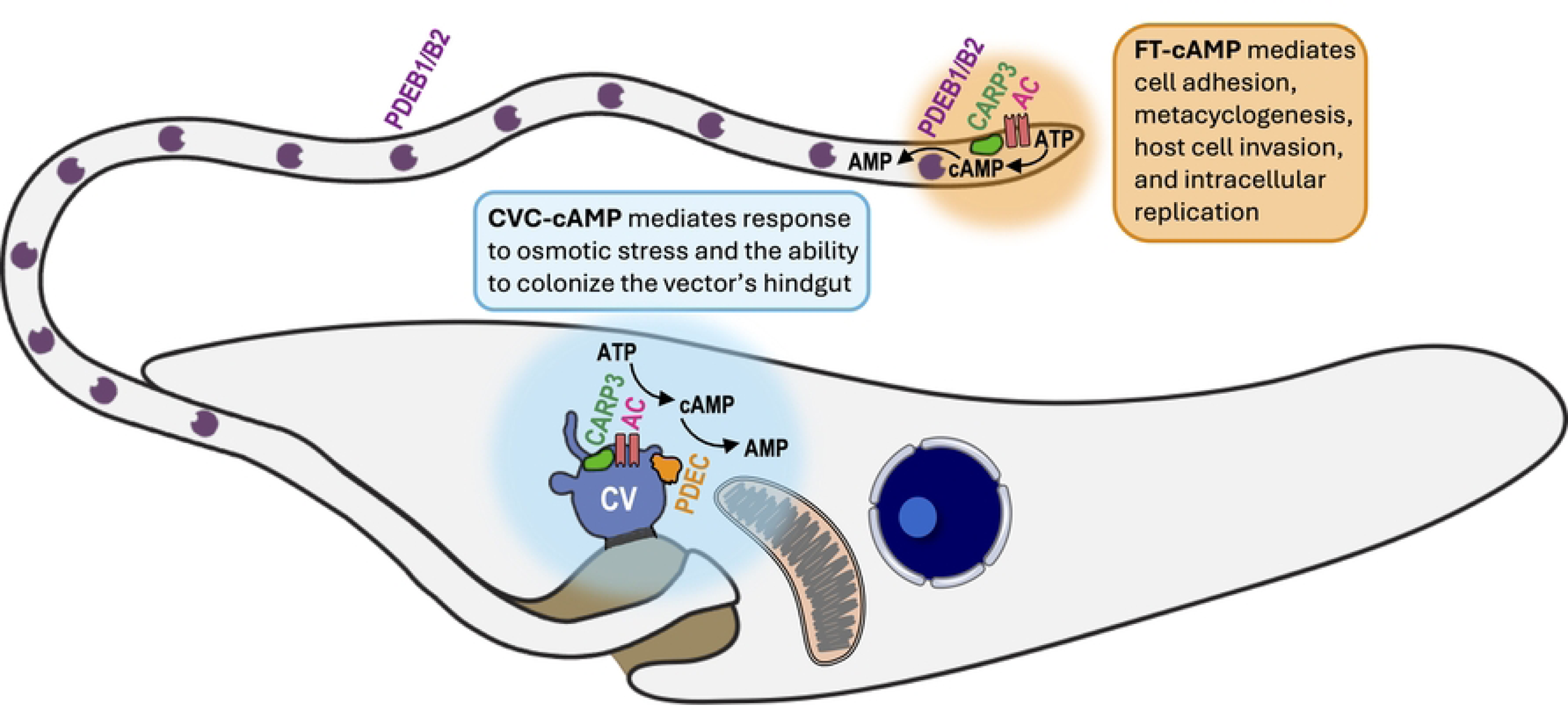
Proposed model for functional segregation of cAMP microdomains in *T. cruzi*. Osmotic stress activates an adenylate cyclase (AC) modulated by cAMP response protein C (CARP3) at the central vacuole (CV) of the contractile vacuole complex (CVC), leading to the synthesis of a localized cAMP pool compartmentalized and maintained by phosphodiesterase C (PDEC). CVC-cAMP signals (indicated in blue) specifically drive osmoregulatory processes that are necessary for parasite survival within the vector’s gut. Conversely, different external cues, such as nutrient depletion or parasite contact with vector’s tissues or host cells, stimulate an AC modulated by CARP3 to synthesize a separate cAMP pool at the flagellar tip (FT), compartmentalized and maintained by phosphodiesterases B1 and B2 (PDEB1 and PDEB2) localized along the flagellum. FT-cAMP signals (indicated in orange) mediate epimastigote adhesion, metacyclogenesis, invasion of mammalian cells by cell-derived trypomastigotes, and intracellular replication of amastigotes. Collectively, our findings support a model in which PDEs determine the spatial and functional separation of CVC-cAMP and FT-cAMP pools, mediating specific cellular responses to environmental changes encountered by *T. cruzi* during its life cycle.

In conclusion, we demonstrate that compartment-specific PDEs regulate distinct cAMP microdomains in *T. cruzi*, orchestrating critical physiological processes required for parasite response to environmental stress, differentiation, infectivity, and ultimately transmission. The different roles of FT-cAMP and CVC-cAMP unequivocally demonstrate the importance of compartmentalized cAMP signals as drivers of signal specificity. Targeting these PDE-regulated pathways represents a promising approach for interrupting *T. cruzi* transmission and ultimately controlling Chagas disease.

## MATERIALS AND METHODS

### Chemicals and reagents

Fetal bovine serum (FBS) was purchased from R&D Systems (Minneapolis, MN). G418 was obtained from KSE Scientific (Durham, NC). Puromycin, blasticidin S HCl, Subcloning Efficiency DH5α competent cells, BCA Protein Assay Kit, SuperSignal West Pico Chemiluminescent Substrate, Horseradish peroxidase (HRP)-conjugated anti-mouse and anti-rabbit IgG antibodies, mouse anti-Ty1 monoclonal antibody, and mouse anti-HA monoclonal antibody, were purchased from Thermo Fisher Scientific (Waltham, MA). Alexa Fluor 488-conjugated donkey anti-mouse and Alexa Fluor 594-conjugated donkey anti-rabbit were from Jackson ImmunoResearch (West Grove, PA). Restriction enzymes and Q5® High-Fidelity DNA Polymerase were obtained from New England BioLabs (Ipsich, MA). ZymoPURE Plasmid Miniprep, ZymoPURE II Plasmid Midiprep, and DNA Clean & Concentrator-5 were from Zymo Research (Irvine, CA). cAMP-Glo™ Assay kit, T4 DNA Ligase, and GoTaq G2 Flexi DNA Polymerase were from Promega (Madison, WI). 4-mm electroporation cuvettes, Precision Plus Protein Dual Color Standards, and nitrocellulose membranes were from Bio-Rad (Hercules, CA). Mouse anti-c-Myc monoclonal antibody (9E10) was from Santa Cruz Biotechnology (Dallas, TX). Fluoromount-G mounting medium was from Southern Biotech (Birmingham, AL). The pMOTag23M vector [88] was a gift from Dr. Thomas Seebeck (University of Bern, Bern, Switzerland). DNA oligonucleotides were purchased from Integrated DNA Technologies (Coralville, IA). Phenylmethylsulfonyl fluoride (PMSF), N-p -tosyl-L-phenylalanine chloromethyl ketone (TPCK), trans-epoxysuccinyl-l-leucylamido-(4-guanidino) butane (E64), protease inhibitor cocktail for use with mammalian cell and tissue extracts, Benzonase nuclease, and all other reagents of analytical grade were from Sigma-Aldrich (St. Louis, MO). Adult *Rhodnius prolixus*, Strain CDC, NR-44077, was provided by the Centers for Disease Control and Prevention for distribution by BEI Resources, NIAID, NIH.

### In silico analyses

The nucleotide sequences of genes encoding *T. cruzi* (Y strain) phosphodiesterases used in this study were retrieved from TriTrypDB [89], a kinetoplastid informatics resource database (tritrypdb.org): *TcPDEA* (TcYC6_0116440), *TcPDEB1* (TcYC6_0028510), *TcPDEB2* (TcYC6_0028500), *TcPDEC* (TcYC6_0111120), and *TcPDED* (TcYC6_0110850). Phylogenetic analyses were conducted in MEGA 12 using the Neighbor-Joining method [90] and the bootstrap method with 1,000 replicates [91]. The phylogenetic tree distances were computed using the JTT matrix-based method [92]. The rate variation among sites was modeled with a gamma distribution (shape parameter = 4). The Gene IDs of all PDE sequences included in the phylogenetic tree are listed in Table S1. The protein domains of TcPDEA, TcPDEB1, TcPDEB2, TcPDEC, and TcPDED were predicted using the InterPro database (https://www.ebi.ac.uk/interpro/) [93] and verified in TriTrypDB. The selection of protospacers for CRISPR/Cas9 gene KO and tagging was performed using EuPaGDT (eukaryotic pathogen CRISPR guide RNA/DNA design tool; http://grna.ctegd.uga.edu) [94]. Sequence analysis and primer design were performed using Benchling (https://benchling.com/).

### Cell culture

*T. cruzi* epimastigotes were grown at 28°C in liver infusion tryptose (LIT) medium supplemented with 10% heat-inactivated fetal bovine serum (FBS), penicillin (100 IU/mL), and streptomycin (100 µg/mL) [95]. The T7/Cas9, pTREX-n-3xHA (EV), and overexpressing cell lines (TcPDEB1-3xHA, TcPDEB2-3xHA, and TcPDEC-3xHA) were grown in 250 µg/mL G418. TcPDEC-3xc-Myc, TcPDEA-3xc-Myc, and TcPDED-3xc-Myc cell lines were maintained with 250 µg/mL G418 and 5 µg/mL puromycin. TcPDEB1/2-KO and TcPDEC-KO cell lines were grown with 250 µg/mL G418, 5 µg/mL puromycin, and 10 µg/mL blasticidin. The growth rate of the OE, KO, and AB epimastigotes was determined by counting cells in culture for at least ten days using a Guava® Muse® Cell Analyzer. Tissue culture-derived trypomastigotes were collected from the culture medium of infected human foreskin fibroblasts (hFFs). hFFs were grown in Dulbecco’s Modified Eagle Medium (DMEM) supplemented with 10% FBS and maintained at 37°C with 5% CO_2_.

### CRISPR-Cas9-mediated gene deletion

To generate null mutant cell lines for *TcPDEC* and *TcPDEB1/2*, *T. cruzi* epimastigotes constitutively expressing T7RNAP and Cas9 (T7/Cas9) were co-transfected with a sgRNA template, and puromycin and blasticidin donor DNAs that served as resistance markers as previously described [62,63]. sgRNA and donor DNAs were amplified by PCR (primers 1-4 for TcPDEC-KO and primers 1,9-11 for TcPDEB1/2-KO, Table S2). Transfected parasites were cultured for 2–3 weeks using G418, puromycin, and blasticidin to select double KO parasites. Gene disruption was verified by PCR using gDNA extracted from mutant parasites (primers 5-8 for TcPDEC-KO and primers 12-17 for TcPDEB1/2-KO, Table S2).

### Generation of overexpression and addback cell lines

We amplified the open reading frames of *TcPDEB1* (using primers 18-19 from Table S2), *TcPDEB2* (primers 20-21 from Table S2), and *TcPDEC* (using primers 22-23 from Table S2) by PCR. Then, we cloned these PCR products into the pTREX-n-3xHA vector by restriction sites XbaI/XhoI [64]. Gene cloning was confirmed by Sanger sequencing, and constructs were used to transfect *T. cruzi* epimastigotes for overexpression of TcPDEs (OE). For the generation of addback cell lines (AB), we cloned the PCR products into the pTREX-h-2xTy1 vector by restriction sites XbaI/XhoI before transfecting the constructs into the corresponding KO cell lines. OE and AB of TcPDEB1, TcPDEB2, and TcPDEC were confirmed by western blot analysis using anti-HA antibodies.

### Endogenous C-terminal protein tagging

As previously described [62,63], the sgRNA templates were designed to perform CRISPR/Cas9-mediated endogenous C-terminal tagging of TcPDEC, TcPDEA, and TcPDED. A donor DNA cassette containing the 3xc-Myc tag sequence and the puromycin resistance gene to induce HDR was amplified using the pMOTag23M vector as a template. The primers used to amplify the donor DNA cassette through PCR were designed to be 60 nucleotides long. These primers included common nucleotide sequences that anneal on the pMOTag23M plasmid and unique target-specific sequences in the forward and reverse primers (Fw/Rv-CTag). The forward primer had a target-specific sequence of 39 nucleotides, while the reverse primer had 34 nucleotides (primers 1,24-26 for TcPDEC, primers 1,29-31 for TcPDEA, and Primers 1, 34-36 for TcPDED, Table S2). *T. cruzi* T7/Cas9 epimastigotes were co-transfected with the sgRNA template and the donor DNA and cultured for 2 weeks with G418 and puromycin to select resistant parasites. Endogenous gene tagging was verified by PCR (primers 27,28 for TcPDEC, primers 32,33 for TcPDEA, and primers 37,38 for TcPDED, Table S2) from gDNA and by Western blot analysis using total protein extracts and anti-c-Myc antibodies.

### Transfection of *T. cruzi* epimastigotes

*Trypanosoma cruzi* Y strain epimastigotes were transfected as previously described [62,63]. Briefly, 4 × 10^7^ cells in the early exponential phase were washed with PBS pH 7.4 at room temperature (RT) and resuspended in ice-cold CytoMix (120 mM KCl, 0.15 mM CaCl_2_, 10 mM K_2_HPO4, 25 mM HEPES, 2 mM EDTA, 5 mM MgCl_2_, pH 7.6) at a final density of 1 × 10^8^ cells/mL. Then, 400 µL of cell suspension was transferred to an ice-cold 4 mm electroporation cuvette containing 25 µg of each DNA fragment (purified plasmid or PCR product) in a maximum DNA volume of 40 μL. Three electric pulses (1,500 V, 25 µF) were applied to the cells in cuvettes using a Gene Pulser Xcell Electroporation System (Bio-Rad). Transfected epimastigotes were cultured in LIT medium supplemented with 20% heat-inactivated FBS and the corresponding antibiotics to select resistant parasites until stable cell lines were obtained (2–3 weeks). Clonal populations of transfectant parasites were obtained by serial dilutions in 96-well plates.

### Western blot analyses

Western blots were performed as previously described [96]. Briefly, parasites in the exponential phase of growth were washed in PBS. Then, the parasites were resuspended in radio-immunoprecipitation assay (RIPA) buffer (150 mM NaCl, 20 mM Tris-HCl, pH 7.5, 1 mM EDTA, 1% SDS, 0.1% Triton X-100) plus a mammalian cell protease inhibitor cocktail (1:250 dilution), 1 mM phenylmethylsulfonyl fluoride, 2.5 mM tosyl phenylalanyl chloromethyl ketone, 100 M N-(trans-epoxysuccinyl)-L-leucine 4-guanidinobutylamide (E64), and benzonase nuclease (25 U/mL culture). Cells were incubated for 30 minutes on ice, and protein concentration was determined by BCA protein assay. Thirty micrograms of protein from each cell lysate were mixed with 4x Laemmli sample buffer (Bio-Rad) supplemented with 10% β-mercaptoethanol before application to 8% SDS-polyacrylamide gels. Electrophoresed proteins were transferred onto nitrocellulose membranes with a Trans-Blot Turbo Transfer System (Bio-Rad). Membranes were blocked with 5% nonfat dry milk in PBS-T (PBS containing 0.1% Tween 20) overnight at 4°C. Next, blots were incubated for 1 hour at RT with the primary antibody: monoclonal anti-HA (1:2,000), monoclonal anti-c-Myc (1:1,000), or monoclonal anti-Ty1. After three washes with PBS-T, blots were incubated with the secondary HRP-conjugated antibody (goat anti-mouse IgG or goat anti-rabbit IgG, 1:10,000 dilution). Membranes were washed three times with PBS-T and incubated with Pierce ECL Western Blotting Substrate (Thermo Fisher Scientific). Finally, images were acquired and processed with a ChemiDoc Imaging System (Bio-Rad).

### Immunofluorescence assay

IFA was performed as previously described [27]. Briefly, *T. cruzi* (epimastigotes, metacyclic trypomastigotes, trypomastigotes, or amastigotes) were washed with PBS and fixed with 4% paraformaldehyde (PFA) in PBS pH 7.4 for 1 hour at RT. IFAs involving TcPDEC mutants were performed under hypoosmotic conditions. Then, cells were allowed to adhere to poly-L-lysine-coated coverslips and permeabilized for 5 minutes with 0.1% Triton X-100. Cells were blocked with trypanosome blocking solution (3% bovine serum albumin [BSA], 1% fish gelatin, 5% normal goat serum, and 50 mM NH_4_Cl in PBS pH 7.4) overnight at 4°C. Cells were incubated with primary antibody(es) mouse anti-c-Myc (1:100), mouse anti-HA (1;200), mouse anti-Ty1 (1:200) diluted in 1% BSA in PBS (pH 8.0) for 1 hour at RT. After washing three times with 1% BSA in PBS, cells were incubated for 1 hour at RT with Alexa Fluor 488-conjugated goat-anti mouse (1:1,000). Then, cells were washed three times with 1% BSA in PBS and mounted on slides using Fluoromount-G mounting medium containing 5 µg/mL 4,6-diamidino-2-phenylindole (DAPI) to stain DNA. Differential interference contrast (DIC) and fluorescence optical images were captured using a Nikon Ni-E epifluorescence microscope on 100× oil immersion lens using NIS-Elements software for acquisition and subsequent processing of the images.

### Determination of intracellular cAMP

The luminescent-based assay cAMP-Glo (Promega) was used to determine the intracellular levels of cAMP in *T. cruzi* epimastigotes [28]. *T. cruzi* epimastigotes in the exponential phase of growth were washed twice with PBS and suspended in an induction buffer (500 μM 3-isobutyl-1-methylxanthine and 100 μM Ro 20-1724 in PBS, pH 7.4) to a final density of 1 × 10^9^ cells/mL. Then, 10 μL of the cell suspension was transferred into a white 96-well plate in triplicates (1 × 10^7^ cells/well). The cells in the wells were lysed by adding 10 μL of cAMP-Glo lysis buffer and incubated at room temperature for 15 minutes. Next, 20 μL of cAMP detection solution was added to each well. The cells in the plate were agitated for 1 minute on an orbital shaker and incubated for 20 minutes at room temperature. Finally, 40 μL of Kinase-Glo Reagent was simultaneously added to the wells. After shaking for 1 minute, the plate was incubated for 10 minutes at room temperature. Luminescence was measured using a BioTek Synergy H1 plate reader (Agilent Technologies, Santa Clara, CA, USA). The results from three independent experiments were expressed as mean values of cAMP content relative to control cells.

### RVD and RVI assays

The regulatory volume decrease (RVD) and regulatory volume increase (RVI) after hypo or hyperosmotic stress was monitored as previously described [35]. Briefly, *T. cruzi* epimastigotes in exponential growth were centrifuged at 1,000 × *g* for 7 minutes, washed twice in PBS, and resuspended in an isotonic buffer (64 mM NaCl, 4 mM KCl, 1.8 mM CaCl_2_, 0.53 mM MgCl_2_, 5.5 mM glucose, 150 mM D-mannitol, 5 mM HEPES-Na, pH 7.4, 282 mOsmol/L) at a cell density of 1 × 10^8^ cells/mL. Then, 100 μL aliquots were placed in a 96-well plate in triplicate, and the absorbance was measured every 10 seconds for 3 minutes. After this, the cells were subjected to hypoosmotic or hyperosmotic stress by adding 200 μL hypotonic buffer (64 mM NaCl, 4 mM KCl, 1.8 mM CaCl_2_, 0.53 mM MgCl_2_, 5.5 mM glucose, and 5 mM HEPES-Na, pH 7.4) or 200 μL hypertonic buffer (64 mM NaCl, 4 mM KCl, 1.8 mM CaCl_2_, 0.53 mM MgCl_2_, 5.5 mM glucose, 1 M D-mannitol, 5 mM HEPES-Na, pH 7.4, 1150 mOsmol/L), for a final osmolarity in each well of 115 mOsmol/L or 860 mOsmol/L, respectively. After inducing osmotic stress, absorbance was measured for an additional 15 minutes. The readings were normalized against the values obtained under isosmotic conditions and converted into a percent volume change using the following equation: (Vf – Vo/Vo) × 100, where Vf is the absorbance value at the experimental time point and Vo is the absorbance mean value obtained under isosmotic conditions. The osmoregulatory capacity was quantified using two parameters: the maximum cell volume change and the final volume recovery after hypoosmotic or hyperosmotic stress.

### *In vitro* metacyclogenesis

Metacyclic trypomastigotes were obtained following the protocol described by [97] with minor modifications. Briefly, *T. cruzi* epimastigotes were cultured for 4 days in LIT medium, washed twice in PBS, resuspended in triatome artificial urine (TAU) medium (190 mM NaCl, 17 mM KCl, 2 mM MgCl2, 2 mM CaCl2, 0.035% sodium bicarbonate, 8 mM phosphate, pH 6.9), and incubated for 2 hours at RT. Then, parasites were incubated horizontally for 96 hours in TAU 3AAG medium (TAU medium supplemented with 10 mM L-proline, 50 mM sodium L-glutamate, 2 mM sodium L-aspartate, and 10 mM glucose) in T75 flasks. For quantification, samples were fixed for 1 hour at RT in 4% PFA in PBS, attached to poly-L-lysine-coated coverslips, and washed three times with PBS. Then, parasites were incubated for 1 hour in 50 mM NH4Cl in PBS, washed three more times in PBS, and mounted onto glass slides with Fluoromount-G containing 15 µg/mL DAPI, which stains the DNA present in the nucleus and the kinetoplast of parasites. Twenty fields/slide were analyzed in an epifluorescence microscope with a 100× objective in three independent experiments. Metacyclic trypomastigotes were distinguished from epimastigotes by the location of the kinetoplast in the cell body (posterior in metacyclic trypomastigotes; between the nucleus and the flagellum in epimastigotes).

### Adhesion assay

During *in vitro* metacyclogenesis, parasites adhere to the culture plate surface within 6 hours of horizontal incubation in the TAU 3AAG medium as previously described [45]. Subsequently, fully differentiated metacyclic trypomastigotes get detached and progressively released into the medium for 96 hours. To quantify the ability of *T. cruzi* epimastigotes to adhere to the 12-well plates during the incubation in TAU 3AAG medium, parasite density in the medium was determined at 2, 4, 6, 24, 48, 72, and 96 hours under a light microscope using a Neubauer chamber. The number of adhered parasites was obtained by subtracting the number of non-adhered parasites from the number of total parasites (1.67 x 10^7^ parasites/well) initially added. The results are expressed as mean values of three independent experiments.

### Host cell invasion and intracellular replication

Host cell invasion and intracellular replication were performed as described previously [28]. hFF cells (5 x 10^5^ cells) were plated onto sterile coverslips in 12-well plates and incubated overnight at 37°C, 5% CO2, in DMEM medium supplemented with 10% fresh FBS. Tissue culture-derived trypomastigotes were centrifuged at 1,700 × *g* for 15 min and left standing in a rack at 37°C for 4 h. Competent trypomastigotes swam up and were separated by transferring the supernatant into another conical tube from cell debris and amastigotes from the pellet. The supernatant was centrifuged at 1,700 × *g* for 15 min. Trypomastigotes from the supernatants of these collections were counted to infect hFFs in the coverslips at a 75 multiplicity of infection. At 4 hours, the infection was stopped by aspirating extracellular parasites and washing with pre-warmed Hank’s balanced salt solution (DHANKS) five times to remove extracellular parasites. Then, 1 mL pre-warmed DMEM + 2% fresh FBS was added and incubated for either 24 hours for the invasion assay or 72 hours for the replication assay. After a final wash with PBS, samples were fixed by transferring the coverslip into a clean 12-well plate containing 1 mL of 4% paraformaldehyde in PBS, pH 7.4. After collecting the replication assay coverslips, the plate was stored at 4°C for at least one more hour (72 h post-infection). After a final wash with PBS, pH 7.4, the coverslips were mounted on glass slides in 15 μg/mL DAPI in Fluoromount G. Finally, the slides were sealed with nail polish and analyzed by fluorescence microscopy using a 100× objective. To quantify invasion, 20 fields/slide were visualized on a Nikon Ni-E epifluorescence microscope, and the number of infected and non-infected cells was counted. To quantify the replication of amastigotes, 60 infected host cells were visualized per assay on a Nikon Ni-E epifluorescence microscope, and the number of amastigotes per infected cell was counted.

### Infection of kissing bugs with *T. cruzi* parasites

Kissing bugs (*Rhodnius prolixus*), were obtained from the colonies established at the Centers for Disease Control (BEI Resources, NR-44077) [98]. These kissing bugs were fed artificially bi-weekly with defibrinated rabbit blood (Hemostat) with a parafilm membrane feeding system (Hemotek). The colony condition was held at 24.0 ± 0.5°C, 50 ± 10% relative humidity, and 6:00 am/6:00 pm light/dark photoperiods. Third instar *R. prolixus* were collected for infection with parasites. *T. cruzi* epimastigotes in exponential phase of growth were washed in 5 mL of 1× PBS pH 7.4 and mixed with defibrinated rabbit blood (complement inactivated at 56 ± 0.5°C for 45 min before the addition of parasites) and offered to triatomines at 37°C through an artificial feeder at a concentration of 1×10^8^ parasites/mL [99–101]. The kissing bugs were held under colony conditions to allow for parasite growth and differentiation. Kissing bugs that did not feed on the infected bloodmeal were removed from assessment. After 30 days, the hindguts were dissected out of the triatomine bugs, emulsified in 100μL of 1× PBS pH 7.4, and examined under the microscope for the presence of parasites in the hindgut to establish the percentage of infected insects. Three groups of 9-15 infected kissing bugs were dissected per *T. cruzi* cell line.

### Statistical analyses

Experiments were repeated at least three times (biological replicates), and results are expressed as means ± s.d. of *n* experiments. Parametric statistical methods (e.g., t-test, *F*-test, ANOVA) will be used for comparing normally distributed data (growth, cAMP level, RVD, metacyclogenesis, adhesion, infection, and invasion), and nonparametric tests (e.g., Mann-Whitney U, Wilcoxon rank-sum, Kruskal-Wallis) for non-normally distributed data (intracellular replication), with appropriate corrections for multiple comparisons. Results are considered significant when *P* < 0.05. The analysis of defective and OE mutants will be performed with appropriate control cell lines (T7/Cas9, EV, and/or AB cell lines). We conducted all statistical analyses using GraphPad Prism 10.2 (GraphPad Software in San Diego, CA, USA).

## ACKNOWLEDGMENTS

Funding for this work was provided by the National Institute of Allergy and Infectious Diseases of the U.S. National Institutes of Health (NIH Grants R00AI137322 and R21AI182544 to N.L.). M.A.C. was supported by the U.S. National Institutes of Health (NIH Grant R21AI178573) and the American Heart Association (AHA Grant 23IPA1054779). J.B.B was supported by the U.S. National Institutes of Health (NIH Grants R01AI148551 and R21AI166633). The funding agencies had no role in the study design, data collection, and interpretation, or the decision to submit the work for publication. Opinions contained in this publication do not reflect the opinions of the funding agencies. We declare that we have no competing financial interests.

## SUPPORTING INFORMATION CAPTIONS

**Figure S1. Additional PCR verification of TcPDEC-KO and TcPDEB1/2-KO cell lines. [A]** Predicted sizes of PCR products amplified from TcPDEC-KO genomic DNA using primer sets indicated by arrows. PCR products to verify TcPDEC-KO were resolved by electrophoresis in 1% agarose gel using the following primer sets: **[B]** primers 7 and 8, and **[C]** primers 5 and 8. **[D]** Predicted sizes of PCR products amplified from TcPDEB1/2-KO genomic DNA using primer sets indicated by arrows. PCR products to verify TcPDEB1/2-KO were resolved by electrophoresis in 1% agarose gel using the following primer sets: **[E]** primers 14 and 15, and **[F]** primers 16 and 13, and **[G]** primers 17 and 13. Primer numbers appear as listed in Table S2.

**Figure S2. Fluorescence microscopy of TcPDEB1-AB, TcPDEB2-AB, TcPDEC-AB, and TcPDEC-OE epimastigotes.** The localization of **[A]** TcPDEB1, **[B]** TcPDEB2, and **[C]** TcPDEC in the corresponding addback cell lines was analyzed by IFA. From left to right: the images show TcPDEB1-2xTy1, TcPDEB2-2xTy1, and TcPDEC-2xTy1 in A, B, and C, respectively (green) merged with DIC (left panel) and DAPI (right panel). **[D]** Immunofluorescence assay of TcPDEC-OE epimastigotes. From left to right: the images show TcPDEC-3xHA (green) merged with DIC (left) and DAPI (right). Western blot analysis confirmed the overexpression of TcPDEC-3xHA (107 kDa) in epimastigotes.

**Figure S3. Phenotype of TcPDEB1-OE, TcPDEB2-OE, and TcPDEC-OE in epimastigote growth, metacyclogenesis, and adhesion *in vitro*.** Growth curve of **[A]** EV, TcPDEB1-OE, TcPDEB2-OE, and TcPDEC-OE epimastigotes in LIT medium. **[B]** Metacyclogenesis *in vitro* of EV, TcPDEB1-OE, TcPDEB2-OE, and TcPDEC-OE. Values are means ± S.D., n=3. One-way ANOVA with Dunnett’s multiple comparisons. **[C]** Adhesion assay with EV, TcPDEB1-OE, TcPDEB2-OE, and TcPDEC-OE epimastigotes. Values are means ± S.D., n=3. Two-way ANOVA with Dunnett’s multiple comparisons.

**Figure S4. TcPDEB1/2-KO impairs the intracellular replication of *T. cruzi* amastigotes.** Representative fluorescence microscopy images of hFFs infected with *T. cruzi* trypomastigotes from **[A]** T7/Cas9 (control), **[B]** TcPDEB1/2-KO, and **[C]** TcPDEB2-AB cell lines. At 72 hours post-infection, cells were fixed and stained with DAPI to visualize host cell nuclei and intracellular amastigotes.

**Table S1.** Gene IDs of PDE orthologs used for the phylogenetic analysis of Figure 1E.

**Table S2.** List of oligonucleotides used in this study.

## Notes

### Competing Interest Statement

The authors have declared no competing interest.

